# Two independent modes of chromosome organization are revealed by cohesin removal

**DOI:** 10.1101/094185

**Authors:** Wibke Schwarzer, Nezar Abdennur, Anton Goloborodko, Aleksandra Pekowska, Geoffrey Fudenberg, Yann Loe-Mie, Nuno A Fonseca, Wolfgang Huber, Christian Haering, Leonid Mirny, Francois Spitz

## Abstract

The three-dimensional organization of chromosomes is tightly related to their biological function ^1^. Both imaging and chromosome conformation capture studies have revealed several layers of organization ^2-4^: segregation into active and inactive compartments at the megabase scale ^5^, and partitioning into domains (TADs) ^6,7^ and associated loops ^8^ at the sub-megabase scale. Yet, it remains unclear how these layers of genome organization form, interact with one another, and contribute to or result from genome activities. TADs seem to have critical roles in regulating gene expression by promoting or preventing interactions between promoters and distant *cis*-acting regulatory elements ^9-14^, and different architectural proteins, including cohesin, have been proposed to play central roles in their formation ^15,16^. However, experimental depletions of these proteins have resulted in marginal changes in chromosome organization ^17-19^. Here, we show that deletion of the cohesin-loading factor, *Nipbl*, leads to loss of chromosome-associated cohesin and results in dramatic genome reorganization. TADs and associated loops vanish globally, even in the absence of transcriptional changes. In contrast, segregation into compartments is preserved and even reinforced. Strikingly, the disappearance of TADs unmasks a finer compartment structure that accurately reflects the underlying epigenetic landscape. These observations demonstrate that the 3D organization of the genome results from the independent action of two distinct mechanisms: 1) cohesin-independent segregation of the genome into fine-scale compartment regions, defined by the underlying chromatin state; and 2) cohes-dependent formation of TADs possibly by the recently proposed loop extrusion mechanism ^20,21^, enabling long-range and target-specific activity of promiscuous enhancers. The interplay between these mechanisms creates an architecture that is more complex than a simple hierarchy of layers and can be central to guiding normal development.

---

Different architectural protein complexes have been proposed to contribute to interphase chromosomal organization, notably the site-specific DNA-binding factor CTCF and cohesin, which co-localize at TAD boundaries and loops ^6,8 22,23^. Mutations of different cohesin subunits in humans lead to different disorders including Cornelia de Lange syndrome ^24^, which are proposed to result in part from gene mis-expression due to defective chromatin folding. While recent models and experimental evidence suggest that regions bound by CTCF and cohesin contribute to organizing loops and TAD boundaries ^8,9,20,25,26,27^, experimental depletion of cohesin subunits or CTCF have shown a limited impact on local chromatin interactions, with most structures, including TADs, appearing largely unaffected ^17-19^. Therefore, it remains unclear to what extent these complexes play a part in the genome’s 3D organization.

Here, instead of depleting cohesin’s constitutive subunits, we interfered with the loading of cohesin complexes by deleting the loading factor *NIPBL/SCC2*. Our reasoning was that since the turnover of chromosome-bound cohesin is relatively fast in interphase (in the range of few minutes) ^28,29^, constant loading is required for its presence on DNA. Efficient deletion of *Nipbl* in non-dividing hepatocytes was achieved by using a liver-specific, tamoxifen-inducible Cre driver (Fig.1a). This conditional, inducible approach allowed us to circumvent the lethality of *Nipbl* +/- mice and the essentiality of cohesin in dividing cells (Extended Data Figure 1). Ten days after tamoxifen injection, *Nipbl* expression was dramatically reduced (Fig.1b) and led to a displacement of cohesin proteins from the chromatin fraction to the soluble nuclear fraction, indicating a strong eviction of cohesin from chromosomes (Fig.1c). No particular pathological signs compared to control animals (either mock-injected *Nipblflox/flox*; *App-Cre* animals or tamoxifen-injected *Nipbl*+/+; *App-Cre* animals) were observed in the liver, and hepatocytes showed no sign of either cell death or proliferation (Extended Data Figure 1).

**Figure 1.**
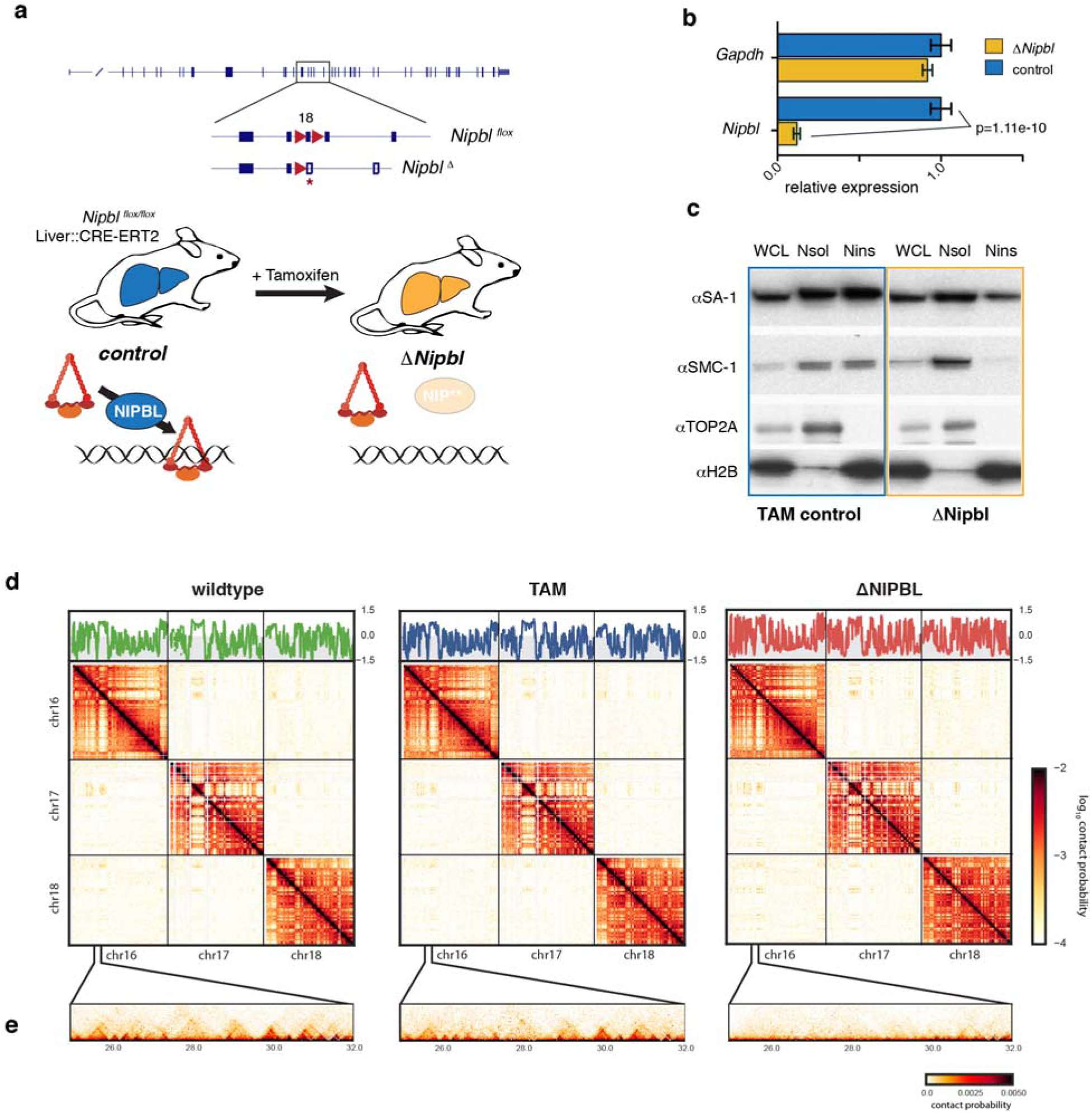
Overview of the experimental design. (a) A conditional *Nipbl* allele was created by flanking exon 18 with *loxP* sites. Upon CRE-mediated recombination, this exon is deleted leading to a frameshift and production of a non-functional protein (Fig. S1). A liver-specific driver for the conditional CRE-ERT2 fusion protein allows precise control of the deletion of *Nipbl,* which occurs in adult hepatocytes only after injection of Tamoxifen. In absence of *Nipbl*, cohesin (represented by a triangular ring) is not loaded on chromatin. (b) Expression level of *Nipbl* and *Gapdh* (control) by RT-qPCR in control (n=4) and ΔNipbl (n=5) hepatocytes. Mean normalized gene expression (using *Pgk1* as internal control, and with expression level in WT set as 1) is displayed as mean and s.e.m. Statistical difference was assessed with unpaired *t*-test.(c) Western blots of hepatocytes protein extracts (WCL: whole cell lysate, Nsol (nuclear extract, soluble fraction) Nins (insoluble, chromatin fraction of nuclear extract)) showed displacement of cohesin structural subunits (SA-1, SMC1) from the chromatin-bound fraction. The efficient separation of the two fractions is shown by the distribution of the TOP2Aand H2B proteins (d-e) Hi-C contact maps at 20kb resolution of WT (left), TAM control (middle) and ΔNipbl cells (right). Top – compartment tracks calculated via eigenvector decomposition of intrachromosomal contact matrices. Middle – *cis* and *trans* contact maps of chr16-18. **(e)** an example of short-range contact patterns in the region chr16:25-32Mb.

To assess the consequence of *Nipbl* depletion and cohesin unloading on chromosome organization, we performed tethered chromatin conformation capture (TCC, referred below as Hi-C) ^30^ on purified hepatocytes from wildtype (WT), tamoxifen control (TAM) and ΔNipbl animals (Fig.1d). For each of these three conditions, two biological replicates were generated. The contact maps obtained from each biological replicate showed extensive similarities (Extended Data Figure 2). For further analyses, we pooled the two replicate datasets to generate Hi-C maps for the three different conditions. In parallel, we determined the complete transcriptome of WT, TAM and ΔNipbl hepatocytes by strand-specific RNA-seq, and performed ChIP-seq for H3K4me3 and H3K27ac to compare the regulatory programs of the different cell populations (**see Supplemental Methods**). We compare Hi-C maps for ΔNipbl and control cells at each characterized level of chromatin organization (Extended Data Figure 3): compartments, TADs, loops, and global scaling of the contact probability *P(s)* with genomic separation *s*^31^.

While previous studies using cohesin depletion observed very mild effects on TADs (Extended Data Figure 4), our Hi-C data reveal a striking effect of *Nipbl* deletion on genome organization (Fig.1d). Compared to WT and TAM control samples, ΔNipbl cells show a genome-wide disappearance of TADs, leaving genome compartmentalization relatively unaffected (Fig.1d). Disappearance of TADs in ΔNipbl is widespread and evident in individual maps (Fig.2a), as well as on the composite map constructed by averaging the ΔNipbl Hi-C map around locations of TAD boundaries detected in WT maps (Fig2b, Extended Data Figure 5). While in regions of inactive chromatin no other structures emerge upon loss of TADs, some local organisation is retained in active and repressed regions (Extended Data Figure 5, Supplemental Methods). As shown below, these structures reflect fine A/B compartmentalization of the genome, rather than residual TADs.

**Figure 2.**
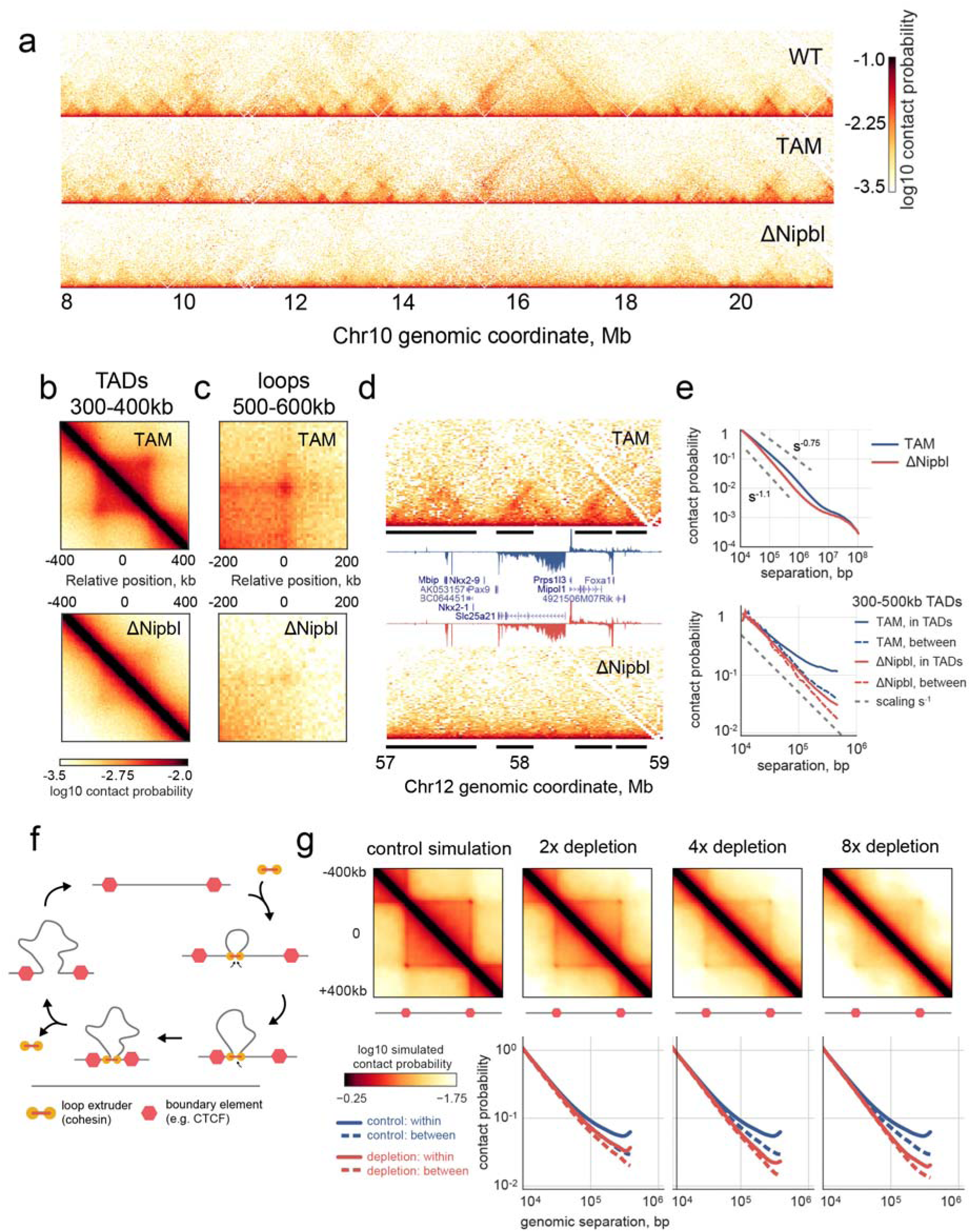
*Nipbl* deletion leads to disappearance of TADs and loops from Hi-C contact maps. **(a)** The short-range contact map for chr10:8-21Mb illustrating robust uniform disappearance of TADs and loops. **(b)** The average map of 564 TADs of length 300-400kb and **(c)** the average map of 102 loops of size 500-600kb. **(d)** The contact map of an example 2Mb region chr12:57-59Mb with overlaid expression track. Top and bottom panels – Hi-C contact maps in TAM control and ΔNipbl, correspondingly, black bars show TADs detected in WT contact maps. Middle panel – location of annotated genes and RPM normalized RNA-seq tracks in sense (above the axis) and anti-sense (below) directions, TAM in blue, ΔNipbl in red. **(e)** Top - genome-wide curves of contact frequency P(s) vs genomic distance s in TAM control and ΔNipbl, normalized to unity at s=10kb. Bottom – P(s) curves plotted separately for contacts formed within or between TADs of size 300-500kb. **(f)** The cartoon representation of the loop extrusion model of cohesin action, which explains how cohesins can form TADs and loops ^21^. In this model, cohesins form cis-loops by first binding to adjacent loci on chromosomes (top and right diagrams). After binding, cohesins translocate along the fiber in both directions, effectively extruding a loop (bottom diagrams). Extrusion halts when cohesins reach boundary elements, formed either by a bound CTCF or any other impediment to extrusion. Extruded loops disassemble when cohesins unbind from the chromosome (left diagram). **(g)** Polymer simulations of loop extrusion reproduce the effects of cohesin depletion on Hi-C contact maps. Top row – average maps of TADs formed on contact maps in polymer simulations of loop extrusion. Left-to-right: the impact of sequential cohesin depletion on the contact map of a TAD in simulations. Bottom row –contact frequency, P(s), versus genomic distance, s, calculated separately for contacts formed within and between TADs.

Loops between specific elements within TADs, which are visible as bright corners in about 50% of TADs ^8^, disappear in ΔNipbl maps too, showing up to 4 fold reduction in contact frequency (Fig.2c and Extended Data Figures 2c and 4). Similarly, the insulation and directionality of the contact footprint of CTCF sites disappear upon *Nipbl* deletion as seen in composite maps centered at oriented CTCF sites (Extended Data Figure 6). Importantly, these structural changes cannot be attributed to altered expression in ΔNipbl cells, as TADs vanished even in regions with unchanged expression (Fig.2d) as well as in regions with up-and down-regulated transcription (Extended Data Figure 5c). This major reorganization of chromatin architecture is also reflected in the curves of contact frequency *P(s)* as a function of genomic distance *s* ^4,32^. In WT or TAM samples, like in other mammalian cells ^20,21^, the scaling of *P(s)* has two different regimes: a more shallow decay for s<200Kb (*P(s)~s^-0.7^*), and a steeper scaling for 200Kb < s < 3Mb (*P(s)~s^-1.2^*). We find that the loss of chromatin-associated cohesin in ΔNipbl cells leads to disappearance of the first regime, producing a single decay of contact probability across the whole range (Fig.2e). This observation suggests that the first scaling regime reflects the compaction of the genome into TADs. We confirmed this by separately calculating *P(s)* within and between TADs: in WT and TAM cells, *P(s)* within TADs decreases more slowly than P(s) for loci separated by a TAD boundary (between TADs). In ΔNipbl, the difference between these two curves is greatly reduced, indicating that the insulating effect of TAD boundaries have diminished and that chromatin folding is more uniform and decondensed (Fig.2e), which is consistent with decondensation observed by imaging upon *Nipbl* reduction ^33^.

TAD formation has been proposed to result from a cohesin-dependent loop extrusion mechanism, where chromatin-associated cohesins extrude progressively expanding loops, until becoming stalled by boundary elements, e.g. bound and properly oriented CTCF ^20,21^ (Fig.2f). Each TAD is a collection of dynamic intra-TAD loops that shows *P(s)~s^-0.7^* intra-TAD scaling. To test whether loop extrusion ^20,21^ can reproduce our experimental findings we simulated the effects of *Nipbl* depletion in a 400kb TAD by reducing the number of loop-extruding cohesins. For each extruding cohesin concentration, we calculated simulated Hi-C maps and *P(s)* within and between TADs. In agreement with our experimental data, 8-fold depletion leads to (i) noticeable disappearance of TADs and loops; (ii) loss of *P(s)~s^-0.7^* regime in the scaling; (iii) decompaction of the chromatin (Fig.2g). Interestingly, TADs were still pronounced at 2-fold cohesin depletion, as has been observed in earlier studies ^17-19^, demonstrating that near-complete removal of chromatin-associated cohesins is necessary for observing a complete disappearance of TADs and loops. Together, these analyses and the observed effects of *Nipbl* deletion indicate that cohesin plays a central role in the compaction of chromosomes into TADs. Moreover, they support the role of cohesin as a loop extruding factor that forms TADs and loops in interphase^20,21^

We next examined the A/B-compartmentalization of chromatin in ΔNipbl cells. Surprisingly, unlike the drastic loss of TADs, compartmentalization still exists in ΔNipbl cells (Extended Data Figure 7) and, in fact, the segregation of A-type and B-type compartmental regions increased (~1.8 fold between the most A- and B-like loci) (Extended Data Figure 8). Closer examination of Hi-C data and compartment tracks, however, reveals widespread local alteration of genome compartmentalization in ΔNipbl cells, with formation of smaller compartmental structures (Fig.3a, Extended Data Figure 9). We found that this persistent compartmentalization of the ΔNipbl genome explains most of the remaining or new domains and boundaries seen in ΔNipbl Hi-C maps (Extended Data Figures 9-10). The finer compartmentalization is reflected in the shorter autocorrelation length of the compartment track (150Kb in ΔNipbl vs ~500Kb in WT and TAM (Extended Data Figure 11), and in the emergence of short B-like compartment regions inside A-regions (Extended Data Figure 11). These emerging B-like regions possess the hallmarks of compartmentalization: (i) they are visible as local depressions in the ΔNipbl compartment track (Fig.3b) and (ii) they show preferential interactions with other B-regions both in far *cis*- and in *trans*-chromosomal maps (Fig.3b). Consequently, they do not represent newly formed TADs since TADs do not exhibit preferential long range interactions. In contrast, B-rich regions, despite a complete loss of TADs, do not experience a similar degree of fragmentation in ΔNipbl cells (Extended Data Figure 11b). Taken together, our observations defy the common notion of TADs simply being the building blocks of larger compartmental segments. Instead, TADs and compartments represent two independent types of chromosomal organization.

**Figure 3.**
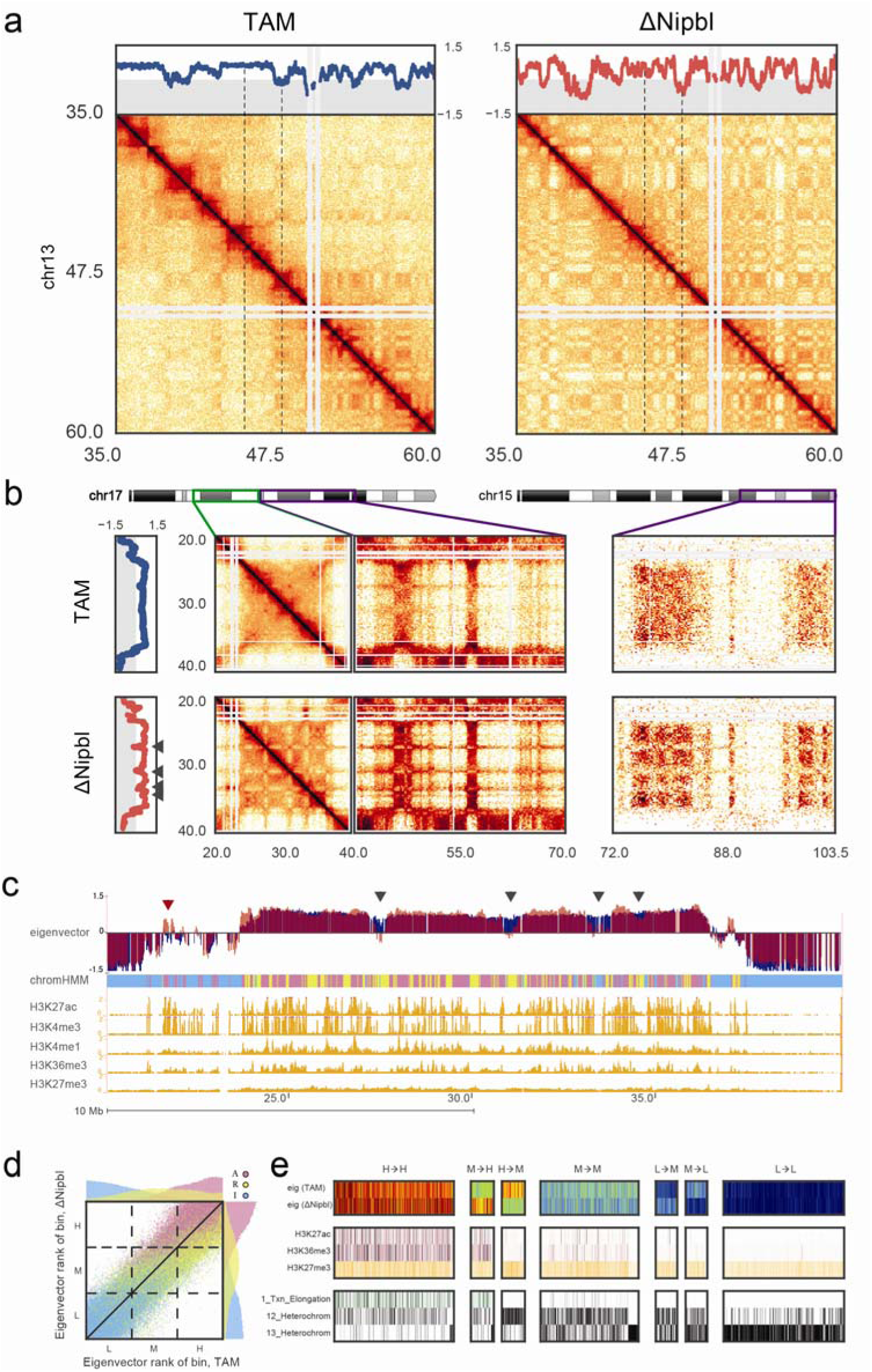
*Nipbl* deletion leads to activity-dependent alteration of compartment structure. **(a)** An example region (chr13:35-60Mb) of a Hi-C contact map showing reorganization of compartment structure. Top panel – compartment signal calculated by eigenvector decomposition of the intra-chromosomal contact matrix at 20kb resolution. Dashed lines indicate the smaller region displayed in Extended Data Figure 11. **(b)** An example of a large region with a uniformly positive (type A) compartment signal in TAM control cells (top) showing compartment fragmentation upon *Nipbl* deletion (bottom). Fragmentation is manifested in the alternating contact patterns of short- (<10Mb, middle left) and long-range *cis* (middle right) and *trans* contact maps (right panel) as well as in the eigenvector compartment signal. Arrows on the left panel indicate the regions of a local drop in compartment signal. **(c)** The loci experiencing a local drop (black arrowheads) in the compartment signal are depleted in epigenetic marks of activity. In contrast, active loci within B-rich regions can show a local increase in compartment signal (red arrowhead). Top to bottom: eigenvector compartment signal in TAM (blue) and ΔNipbl (red) cells; simplified ChromHMM state assignment of loci grouped into active (magenta) / repressed (yellow) / inert (cyan) states; ENCODE activity-related histone ChIP-seq for adult mouse liver cells. **(d)** Rank correlation of 20kb resolution eigenvectors in TAM and ΔNipbl, colored by simplified ChromHMM state. Top and right margins – histograms of eigenvector ranks split by simplified ChromHMM state in ΔNipbl (right) and TAM (top). The dashed lines show the tercile borders, splitting bins into equal-sized groups of low (L), middle (M) and high (H) eigenvector. **(e)** Epigenetic profiles of bins transitioning between eigenvector terciles upon Nibpl deletion. Top to bottom: eigenvector in WT and ΔNipbl cells; ENCODE histone marks; ChromHMM states characteristic of active, repressed and inert chromatin. The bins that transitioned from the middle to the high tercile are enriched in activity marks, while bins transitioning from the high to the middle tercile were depleted in those marks.

Next, we examined how the observed structural reorganization is related to functional characteristics of the genome. Strikingly, we found that the compartment structure of the ΔNipbl genome reflects local transcriptional activity and chromatin state better than the compartment structure of the WT (Fig.3c-e, Extended data Figure 12). The compartment track of ΔNipbl cells shows a stronger correlation with ΔNipbl tracks of activity-associated epigenetic marks, H3K27ac, H3K4me3, and gene expression, as well as related wildtype tracks, e.g. H3K27ac, H3K4me3, DNase hypersensitivity, TF binding etc., smoothed over a wide range of window sizes (Extended Data Figure 11c). To understand the relationship between epigenetic state and the change in compartment status, we compared the compartment tracks to the mouse liver chromatin state segmentation (ChromHMM ^34^) simplified into three state categories: Active, Repressed and Inert (see *Methods*). While the compartmentalization of inert regions is relatively unaffected by *Nipbl* deletion, regions of repressed and active chromatin further diverge in their compartment status (Fig.3cd), producing local peaks in the compartment track in active regions, and local B-like depressions in repressed regions (Fig.3c). Furthermore, regions of facultative lamin-B1 association ^35,36^ are enriched in regions showing a reduction in compartment signal (from A to B-like), while those showing lamin-B1 association across different mouse cell lines are primarily B-type in both WT and the mutant (Extended Data Figure 13, *Methods*). Importantly, these changes in compartmentalization cannot be attributed to changes in expression or in the activity marks (H3K27ac, H3K4me3), which are largely unperturbed in the mutant at the scales relevant to compartmentalization (Extended Data Figures 12,14-15). In summary, in the absence of chromosome-associated cohesin allowd the genome to adopt a compartmental organization that closely reflects the local functional state of the genome. Incidentally, the fact that the association between compartmental structure and epigenetic state is observed even when considering the wildtype epigenetic landscape suggests that the landscape is largely unaffected by loss of chromosome-associated cohesin (Fig.3e, Extended Data Figures 12,15).

Altogether, these results indicate that chromatin has an intrinsic tendency to form small-scale, specific compartments that reflect its local epigenetic landscape and transcriptional activity. However, in wild-type cells, this close association between epigenetic state and 3D organization is reduced, likely because the distinct loop-extruding activity of cohesin that leads to formation of TADs can bring together and mix loci with opposing states.

Deletion of *Nipbl* and the disappearance of TADs led to altered transcription. About a thousand genes are significantly mis-expressed (637 down-regulated, median fold-change=0.32, 487 up-regulated, median fold-change=3.15, with DESeq2 tools) upon *Nipbl* deletion and TAD disappearance (Fig.4a and Extended Data Figure 16). Importantly, while H3K27ac (and H3K4me3) peaks at promoter regions of affected genes change in coherence with expression changes, distal peaks (marking active distant enhancers) are mostly unaffected (Extended Data Figure 17). This conservation of distal enhancer signal indicates that the regulatory potential of the cells was mostly unperturbed, yet transcriptional changes did occur. We noticed that down-regulated genes are surrounded by a larger intergenic space (defined by the distance separating the TSS of their immediate neighbours) than up-regulated or unaffected ones (Extended Data Figure 16b) and transcriptional changes are concentrated within regions that normally form larger TADs. If there is no reliable way to identify *a priori* genes for which distal regulatory interactions are essential, this characteristic genomic context of transcriptional alterations is consistent with defective long-range regulatory interactions in ΔNipbl cells. Remarkably, while we saw more genes down-regulated than up-regulated, we noticed an inverse phenomenon in intergenic regions, with widespread up-regulation of exo-genic (intergenic or antisense intragenic) transcription (Fig.4a). Using a conservative approach (see *Methods*), we found 1107 non-genic transcripts or transcribed regions, which showed at least an 8-fold enhanced transcription in ΔNipbl cells; amongst these, 232 correspond to new non-coding RNAs, which are not detected or barely detected in WT or TAM samples, and for most of them not annotated (Extended Data Figure 16d). The new transcription is often bi-directional, (Fig.4bc, Extended Data Figure 17) and occurs either at pre-existing – but often inactive – promoters (marked by small peaks of H3K4me3) or at enhancers (Fig.4d-e, Extended Data Figure 17). We saw several examples of reciprocal expression changes (i.e. down-regulation of a gene being followed by up-regulation of an adjacent gene or of a new non-coding transcripts) (Fig.4b, Extended Data Figure 17e), but often, new non-coding transcription arises without measurable impact on surrounding genes. While the chromatin profile suggests that enhancers retain their normal activity and therefore regulatory potential, this shift from gene-promoter transcription to intergenic transcription initiated in the vicinity of distal regulatory elements suggests that the deletion of *Nipbl* and resulting absence of TADs and loops impairs enhancer communication: with a reduced range of contact, some enhancers cannot reach their target promoters and transfer their activity on nearby alternative, sometimes cryptic, targets (including themselves).

**Figure 4.**
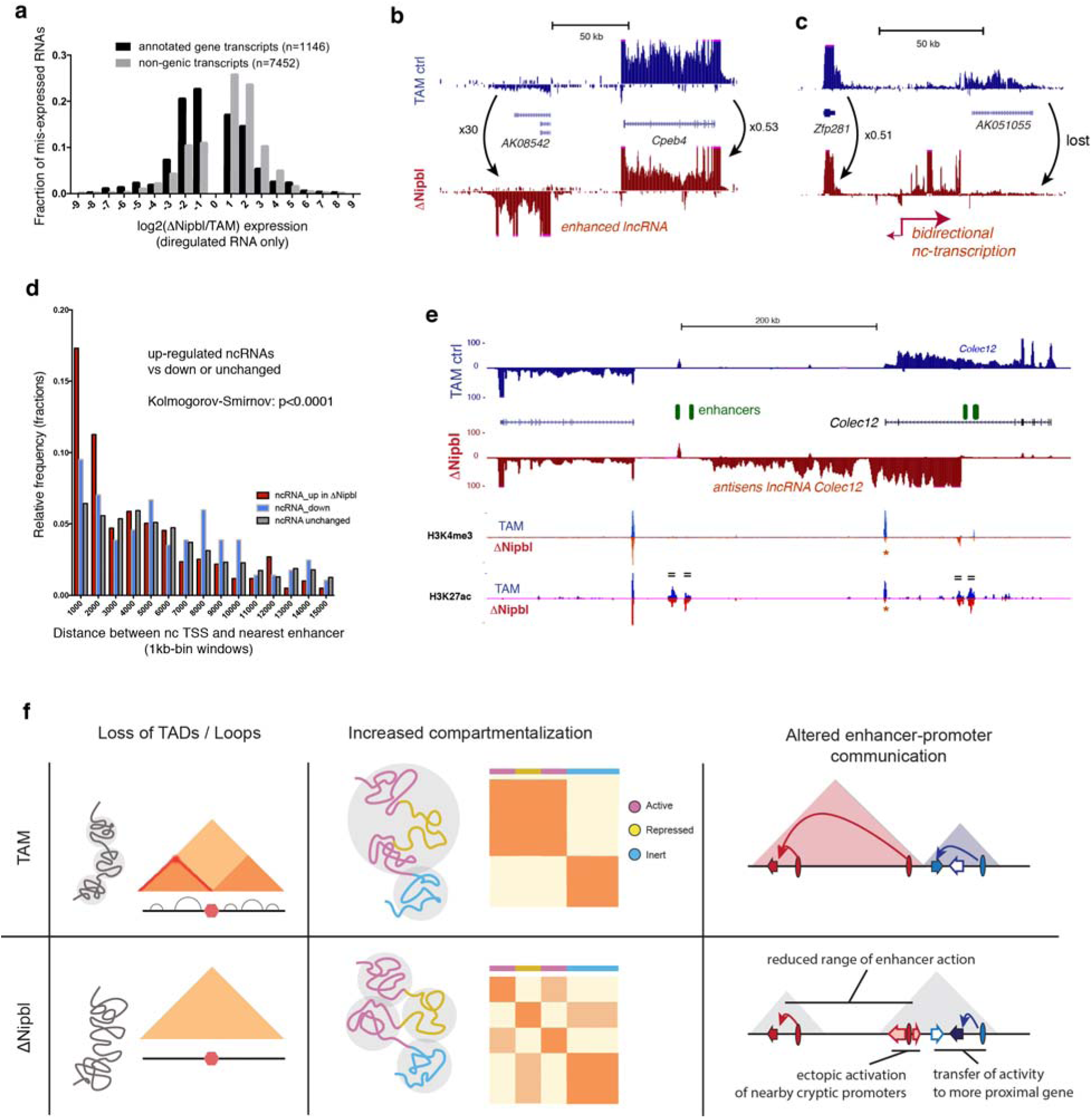
Transcriptional changes in *Nipbl* mutants reveal possible enhancer-promoter communication problems in absence of TADs. **(a)** Distribution of fold-changes for misexpressed genes (black) and exo-genic transcripts (light grey). Genes showing less than a twofold change in either direction were omitted. **(b-c)** Examples of transcriptional changes at different loci, with TAM controls stranded RNA-seq tracks in blue, and ΔNipbl in red. The tracks combined data from four replicates for each condition. Transcriptional changes reported were observed in each replicate. **(d)** Distribution of distances between the transcriptional start of the ncRNAs and the nearest enhancer. **(e)** Transcriptional switch at the *Colec12* locus. RNA-seq tracks show a disappearance of the *Colec12* transcripts, replaced by an antisense transcript initiated from a intronic enhancer. H3K4me3 and H3K27ac profiles show no changes at distal enhancers (green ovals, marked by “=” on the tracks), while peaks at the *Colec12* promoter disappear (red asterisk). **(f)** Summary of the results. TADs (coloured triangles) and loops disappear upon *Nipbl* deletion (left), unmasking a stronger and finer compartmentalization (middle) that is visible as a fragmented plaid pattern in the mutant Hi-C map relative to that of the wildtype and whose alternating member regions more faithfully track transcriptional activity. The resulting reduction of contact range (right) thwarts distant enhancers (ovals) from acting on their normal target genes (arrows, with coloured ones indicating active genes, white ones inactive), leading them to act instead on neighbouring genes or cryptic promoters located in their vicinity. The active units make up new compartmental regions (grey triangle).

Overall, our findings provide new important insights into the mechanisms that organize the genome’s 3D structures and their relation to gene expression. Our data show that local chromatin structures result from the overlapping action of two totally independent mechanisms (Fig.4f). This overlap challenges the standard hierarchical, representation of genome architecture in which individual TADs (or small contact domains) combine to form active (A) or inactive (B) compartmental regions. Partitioning of the genome into these active and inactive compartmental regions and their preferential self-interaction are achieved by a cohesin-independent mechanism. This mechanism acts pervasively, across scales, and notably at a much smaller scale than previously appreciated, reflecting local transcriptional activity and the epigenetic landscape. This fine compartment structure is however blurred by the action of a second, cohesin-dependent mechanism, which compacts chromatin locally, independently of its status, but is constrained by CTCF-enriched boundary elements ^20,21^. This picture is consistent with the formation of dynamic *cis*-loops by a cohesin-dependent loop extrusion process. Our genome-wide data provide support to the notion that cohesin-dependent *cis*-looping and resulting TADs allow for communication between distal regulatory regions in a reliable and tuneable manner ^14^, while the activity-dependent compartmentalization may subsequently reinforce and maintain these interactions.

Mammalian genomic contact maps, revealed by Hi-C, result from the superimposition of these two distinct mechanisms. Their relative contributions can vary between loci, and, for a given locus, between cell types. The co-existence of two processes with different modes and scales of action – together with the intrinsic disorder of chromosome conformations – can explain the difficulties in the field in delineating and unambiguously classifying the different population-averaged patterns seen in Hi-C maps and described as “domains” (TADs^6,7^, subTADs^15^, metaTADs ^37^, physical domains ^38^, contact domains ^8^ etc.), which have led to discussions regarding their existence as structural or functional entities. The experimental ability to now distinguish these two modes of chromosome organization demonstrates that a distinct cohesin-dependent biological mechanism gives rise to the local contact domains identifiable as TADs in Hi-C maps and provides avenues to dissect the process(es) governing their formation and maintenance, as well as to distinguish cause from consequence in the relationship between gene expression and chromatin folding.

## Acknowledgments

This work would not have been possible without the important contribution of the members of the EMBL Laboratory Animal Resources Facility, particularly Silke Feller, for animal welfare and husbandry and of the EMBL Genomics Core Facility for sequencing the different genomic libraries. We thank members of the Mirny and the Spitz labs, John Marioni (EMBL/EBI) and Heather Marlow (IP) for many productive discussions and helpful suggestions. We are grateful to Hugo B. Brandão for providing a MATLAB-based visualization tool. We thank Ana Losada (CNIO) for generously providing antibodies. W.S and A.P were supported by an EMBL Interdisciplinary Postdoc (EIPOD) Fellowship under Marie Curie Actions COFUND. The work in the Mirny lab is supported by R01 GM114190, U54 DK107980 from the National Institute of Health, and 1504942 from the National Science Foundation. The collaboration is also partially supported by MIT-France MISTI Fund. The work in the Spitz lab was supported by the European Molecular Biology Laboratory, the Pasteur Institute and the Deutsche Forschungsgesellschaft (DFG grant: SP 1331/3-1). Funding from the European Commission’s Seventh Framework Programme through the Collaborative Research Project RADIANT (Grant Agreement no: 305626, to W.H.) contributed also to this work.

## Author Contributions

W.S. and F.S. conceived the study and designed the experiments, with input and advice from C.H. W.S. performed all the experiments with the help of A.P for TCC, who also carried out preliminary analyses of the TCC datasets, with advice from W.H.. N.A. and A.G. performed computational analysis of Hi-C, RNA-seq, ChIP-seq and other relevant datasets. Y.L-M and N.A.F. contributed to analysis of transcription data. G.F. performed computer simulations of cohesin depletion and assisted with data analysis and paper writing. L.M. provided advice on data analysis and simulations. W.S., N.A., A.G., L.M. and F.S. wrote the paper with input from the other authors.

## Author Information

The authors declare no competing financial interests. Correspondence and requests for materials should be addressed to F.S. and to L.M.

## EXTENDED DATA

**Extended Data Figure 1.**
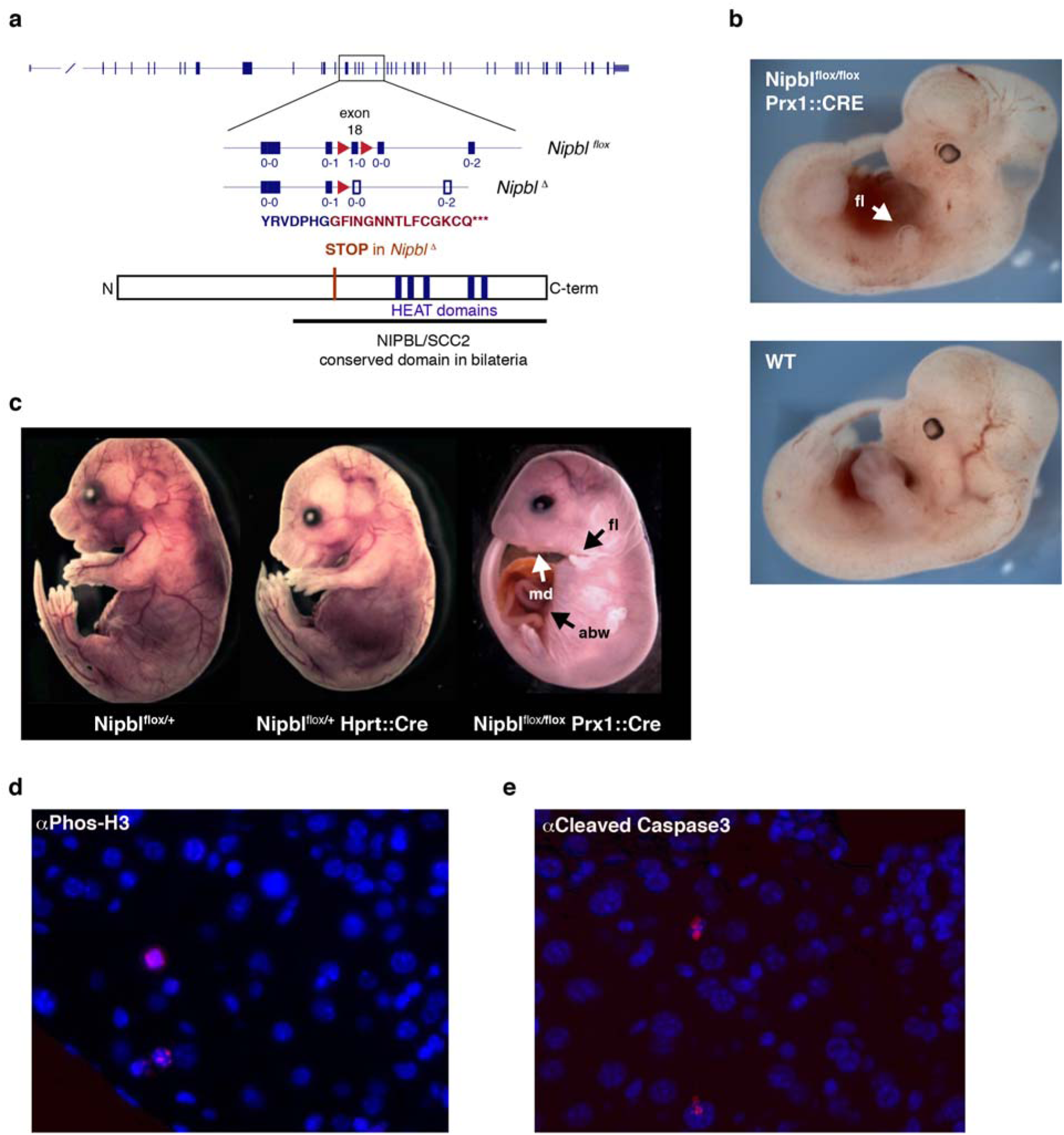
Conditional inactivation of *Nipbl* in mice. **(a)** Schematic representation of the conditional allele, with *loxP* sites (red triangles) flanking exon 18. The reading frame of each exon is indicated below the corresponding square, as “x-x”. Deletion of exon 18 leads to a frame-shift introducing a premature stop codon (indicated by amino acids in red). The resulting protein lacks the critical HEAT domains conserved in NIPBL/SCC2 proteins. **(b-c)** E12 embryos **(b)** and E18 fetuses **(c)** carrying the conditional *Nipbl* allele (*Nipbl^flox^*) and either ubiquitous (Hprt:Cre ^39^) or limb-specific (Prx1::Cre ^40^) Cre recombinase drivers. Structures expressing Cre are rapidly lost in *Nipbl^flox/flox^* animals. Heterozygous *Nipbl^flox/+^* animals are grossly morphologically normal, but die soon after birth, as reported for other *Nipbl* loss of function alleles ^41^. fl=forelimb; md:mandibule; abw=abdominal wall. **(d-e)** Histochemical staining of liver section of adult ΔNipbl hepatocytes (Nipblflox/flox; Ttr::CreERT2; 10 days after Tamoxifen injection) for a proliferation marker (Phos-H3) **(d)** and apoptosis (cleaved Caspase3) **(e)** (both showed in red). Nuclei are stained with DAPI (blue).

**Extended Data Figure 2.**
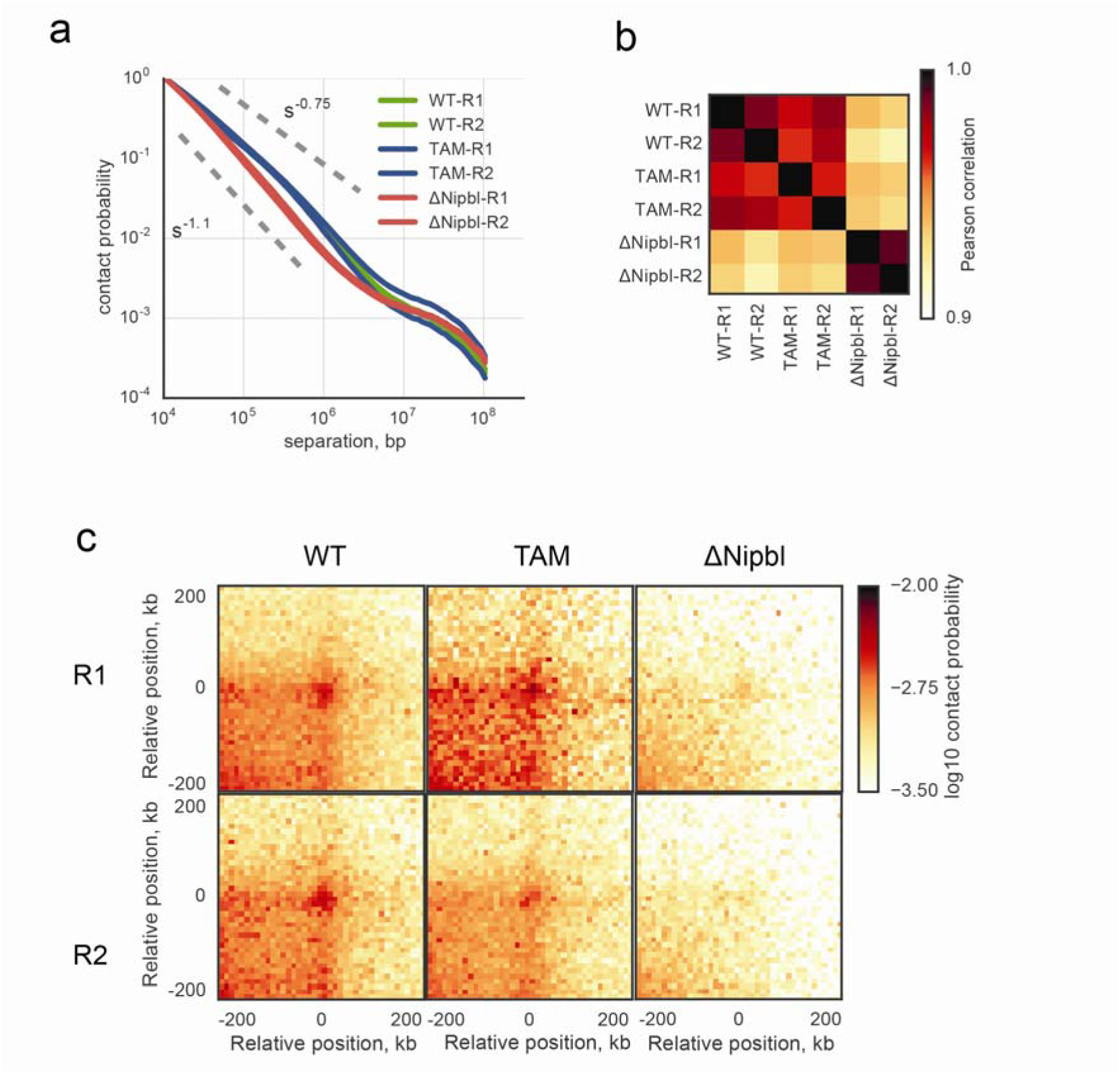
The Hi-C maps obtained in all experimental conditions show extensive similarities between replicates. **(a)** The scaling curves of contact probability P(s) vs genomic distance s, normalized to unity at 10kb separation. **(b)** The matrix of Pearson correlation coefficients between *cis* autosomal eigenvectors of replicates detected at 100kb. **(c)** The average Hi-C maps of 102 500kb-600kb loops ^8^in each replicate of each experimental condition.

**Extended Data Figure 3.**
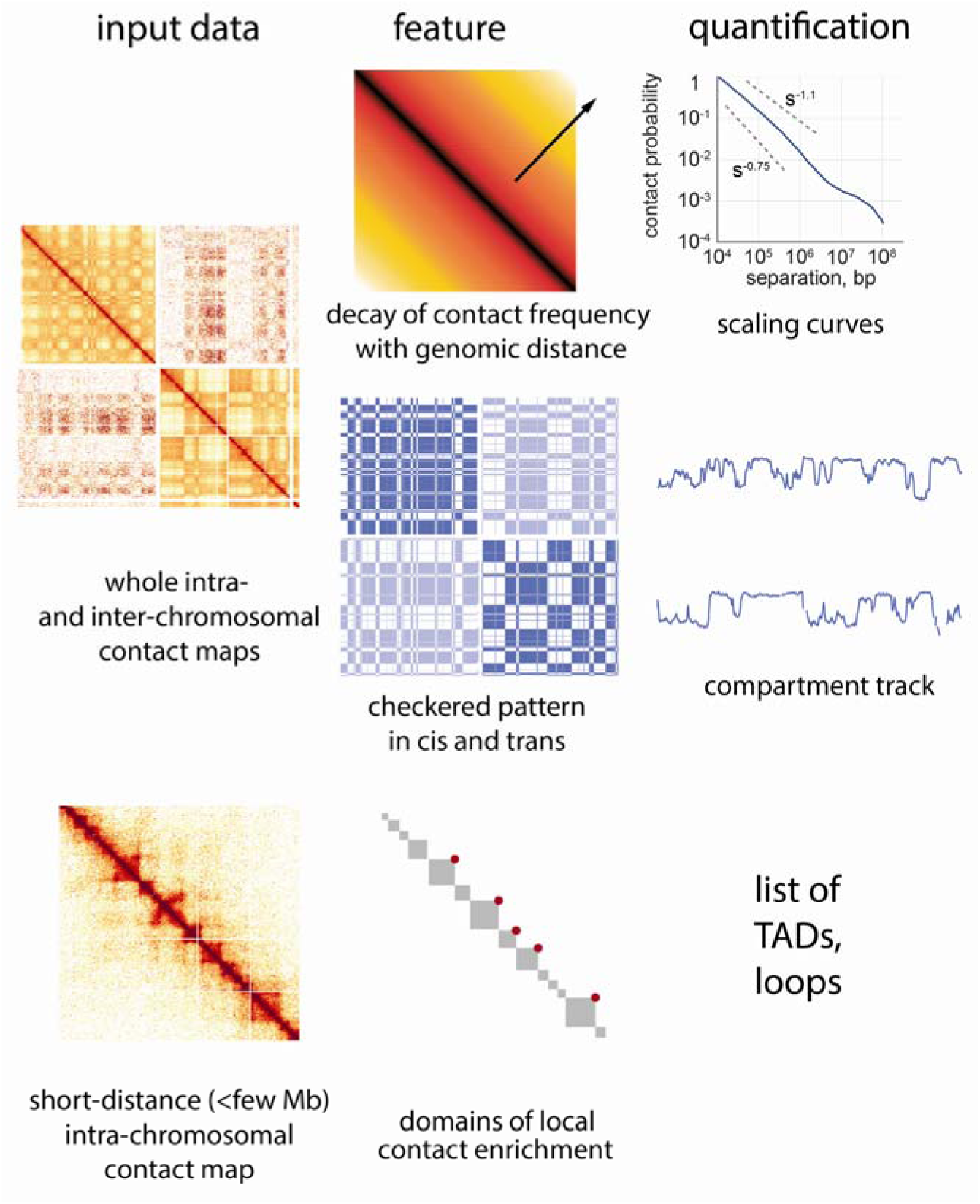
Overview of various features of chromosomal architecture detected and quantified in Hi-C contact maps. Top row – intra-chromosomal maps show the decay of contact frequency with genomic distance, which can be quantified with the curves of contact frequency P(s) vs genomic separation s. Middle row – both intra- and inter-chromosomal maps display a checkered pattern caused by compartmentalization of the genome. This pattern can be quantified by a continuous genomic track obtained via eigenvector decomposition of either *cis* or *trans* maps. Bottom row – intra-chromosomal maps at short genomic distance scales reveal domains of enriched contact frequency, which appear as bright squares along the main diagonal, and loops which appear as bright dots connecting two loci. Both can be detected and quantified using specialized algorithms.

**Extended Data Figure 4.**
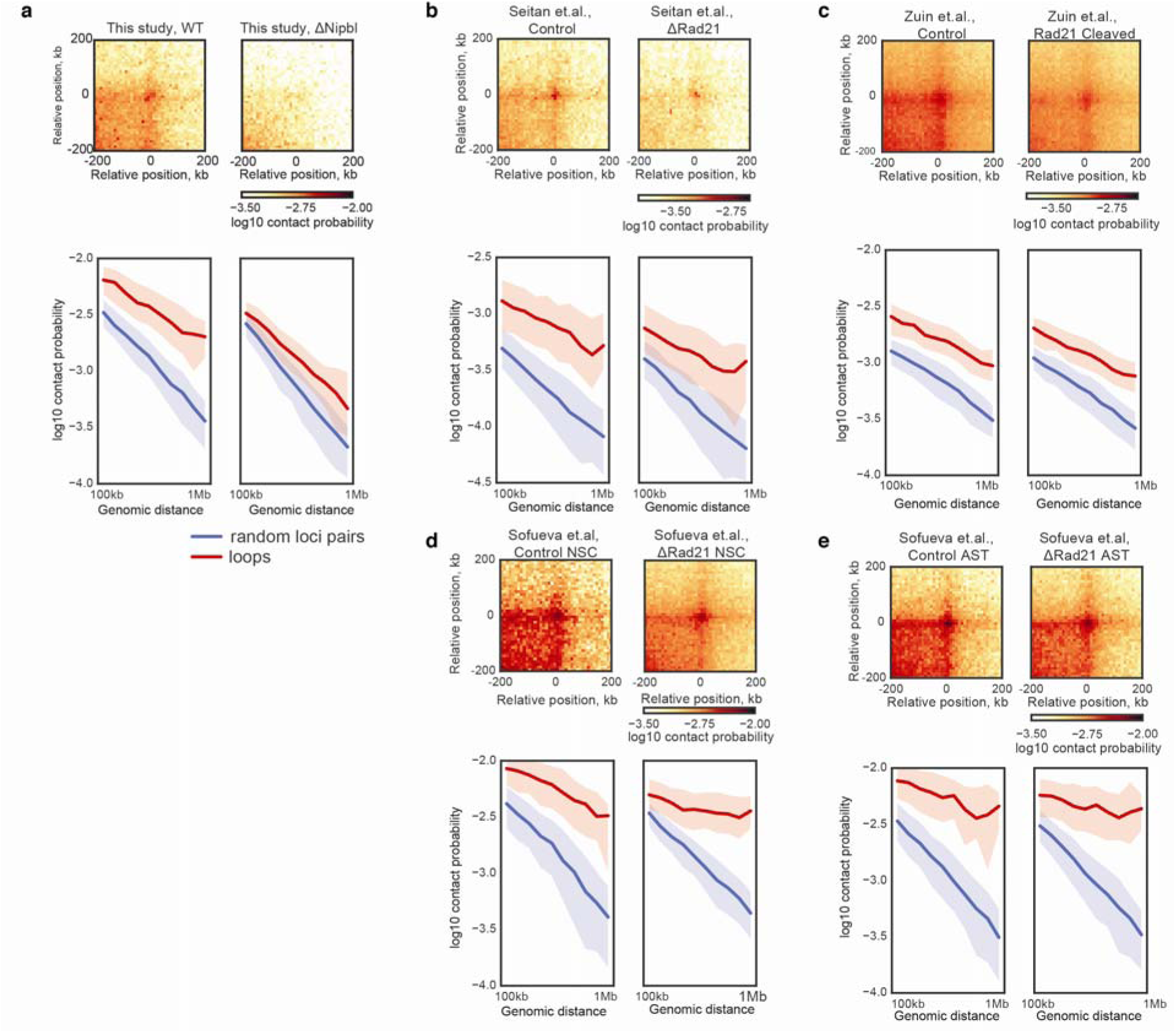
Deletion of *Nipbl* in this study leads to a robust disappearance of loops compared the techniques previously used to deplete chromatin-bound cohesin. **(a)** genetic deletion of *Nipbl* in hepatocytes, this study. **(b)** deletion of *Rad21* in thymocytes ^19^. **(c)** proteolytic cleavage of RAD21 in HEK293T cells ^17^ **(d-e)** deletion of *Rad21* in NSCs and ASTs ^18^. For each dataset: left column, top panels – the average Hi-C map of 102 loops with size range 500-600kb ^8^in WT and ΔNipbl contact maps; bottom panels – the relative contact probability between pairs of loop anchors vs genomic distance, compared to randomly selected pairs of loci. The thick line shows the median contact probability; the shading shows the envelope between the 25^th^ and 75^th^ percentiles of contact probability at each genomic separation.

**Extended Data Figure 5.**
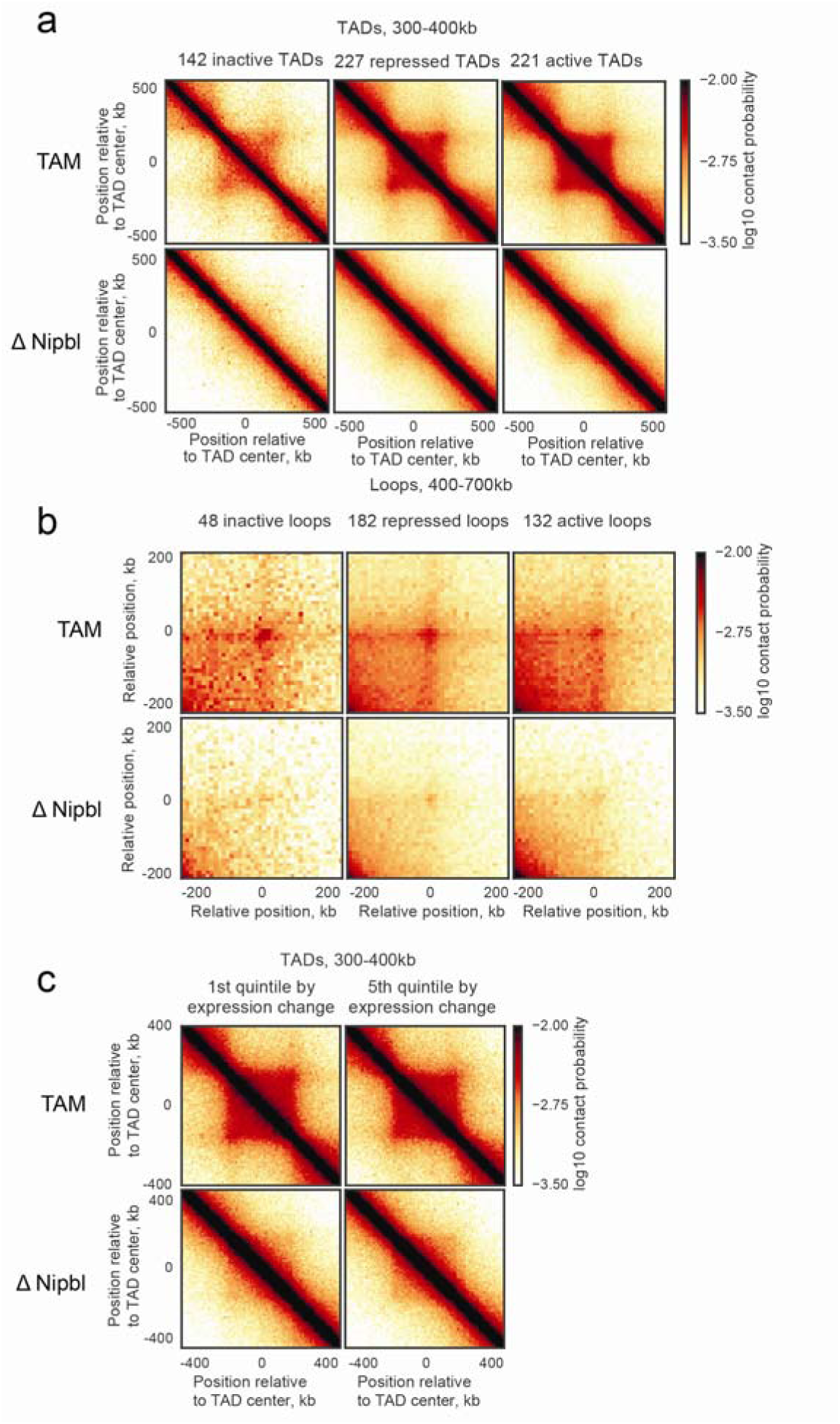
Residual structures are observed in active and repressed regions of the genome after *Nipbl* deletion. For each TAD, an activity was assigned based on the dominant simplified ChromHMM state category. **(a)** The average contact map of 300-400kb TADs in inert, repressed and active regions of the genome. **(b)** The average contact map of 300-700kb loops in inert, repressed and active regions of the genome. **(c)** The average contact maps of most upregulated 20% (left) and most downregulated 20% (right) of 300-400kb TADs.

**Extended Data Figure 6.**
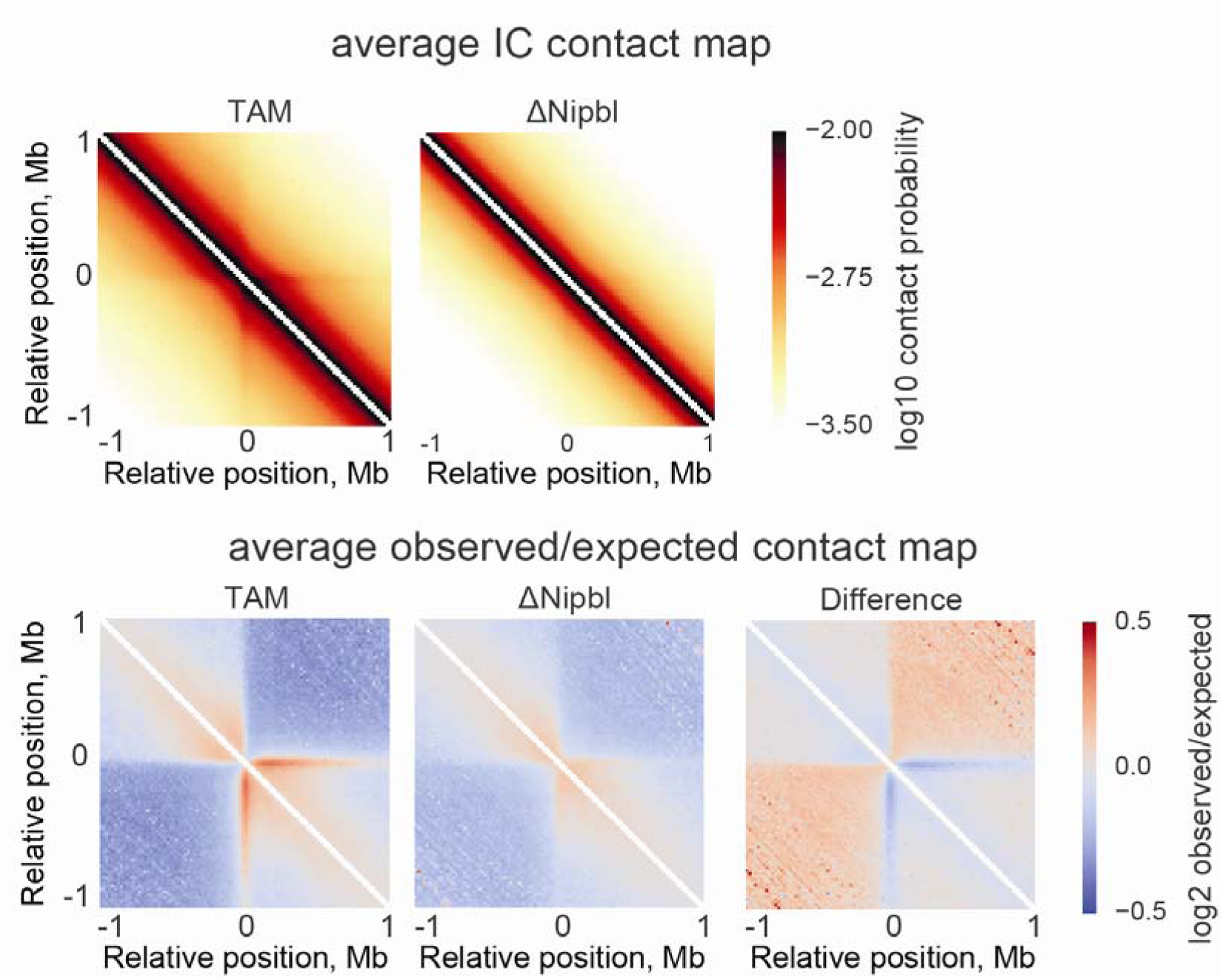
The average Hi-C contact map around CTCF peaks. CTCF peaks are taken from ENCODE mouse liver ChIP-Seq data ^42^and supported by an underlying CTCF binding motif occurrence. Top row – average iteratively corrected contact map around ~22000 sites in TAM and ΔNipbl cells. Bottom row – average contact map normalized by the expected contact frequency at a given genomic separation.

**Extended Data Figure 7.**
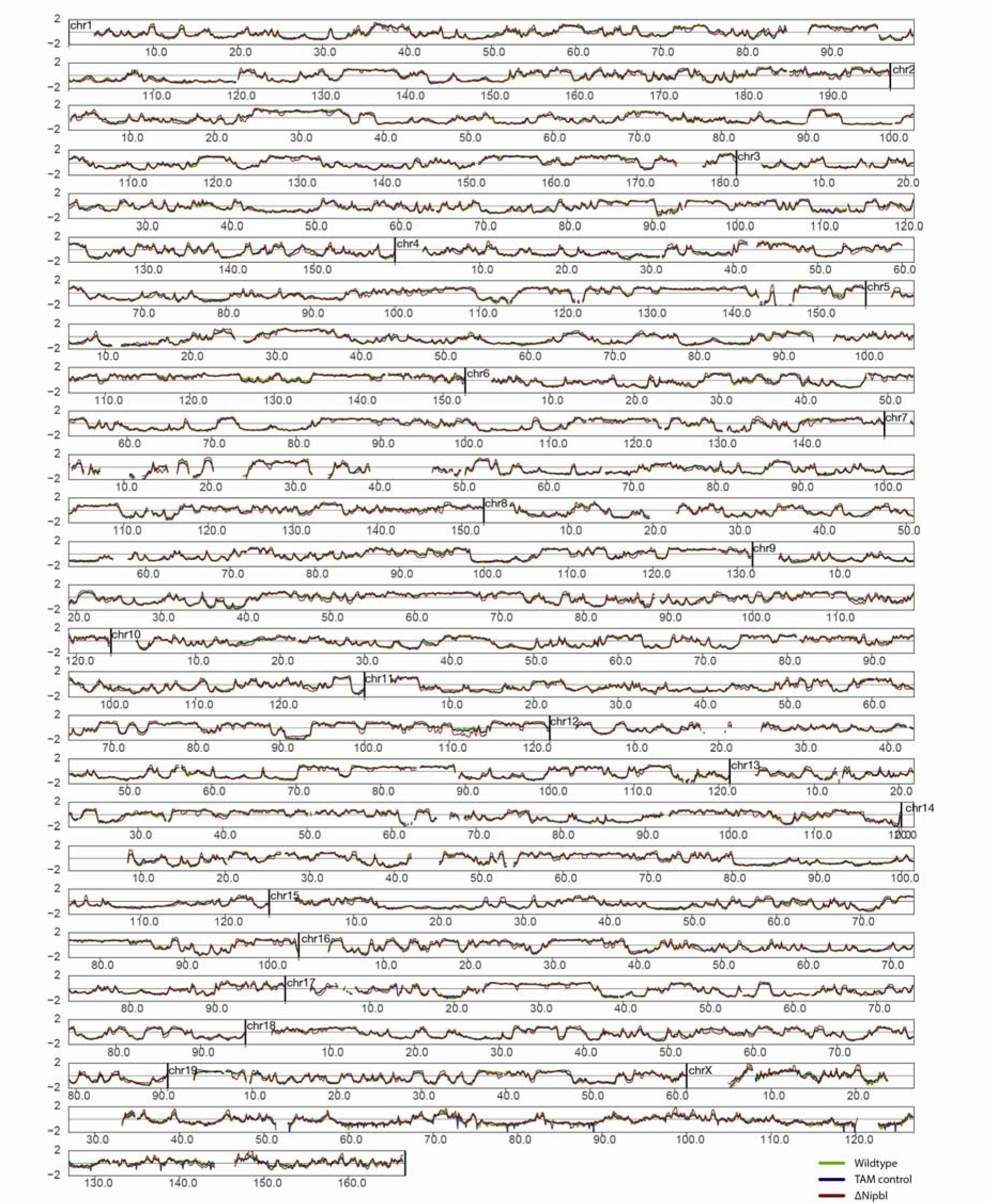
The genome-wide eigenvectors calculated from 20kb *cis* contact matrices in all three experimental conditions, WT (green), TAM (blue), ΔNipbl (red).

**Extended Data Figure 8.**
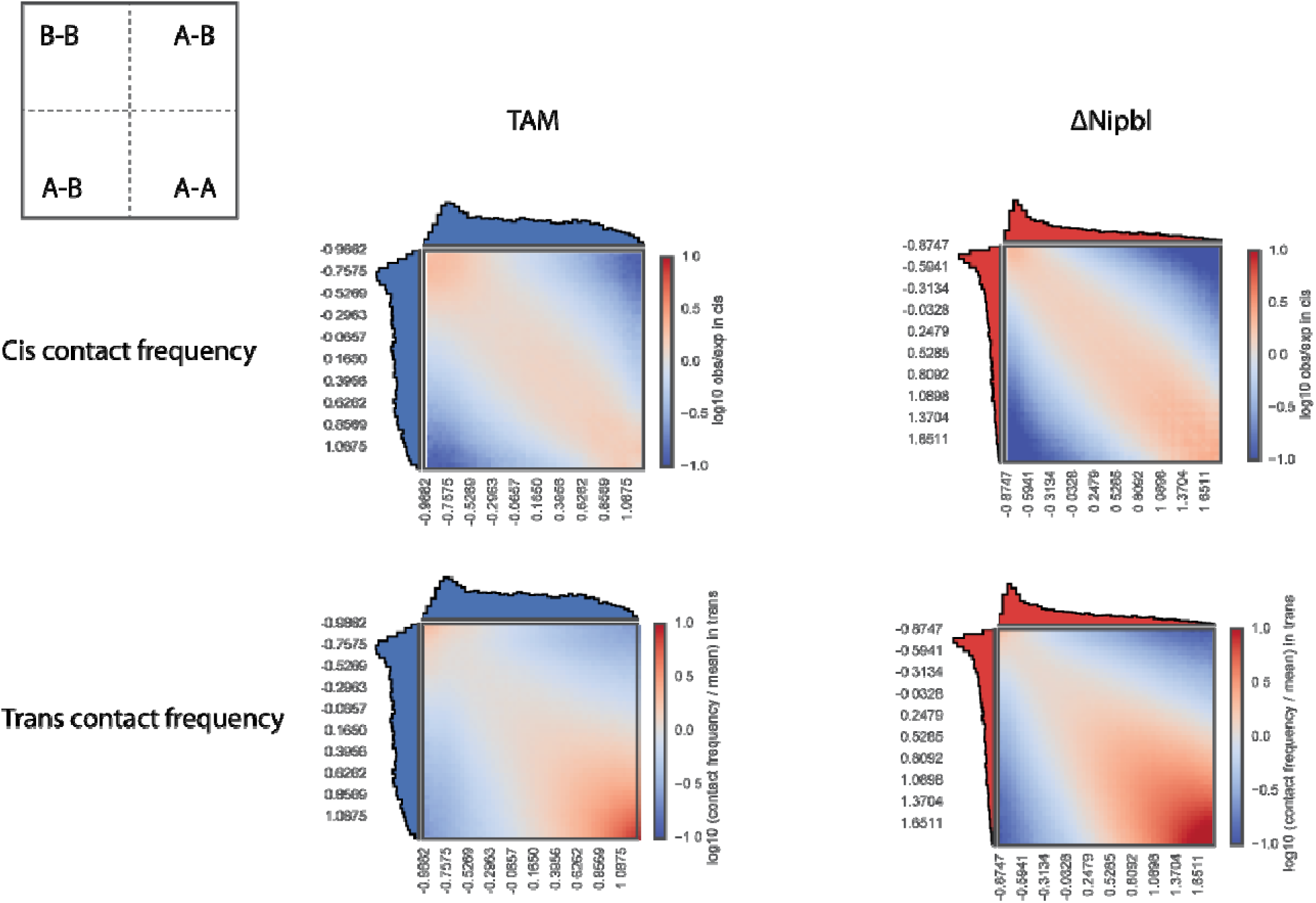
Increased compartmentalisation in ΔNipbl cells. Average interaction frequencies between pairs of loci (100kb bins) arranged by their compartment signal (eigenvector value). Notice enrichment of AA and depletion of AB interactions in ΔNipbl cells: see the diagram for AA, AB, BB regions. The interaction frequencies in *cis* (top row) are computed for observed/expected contact maps. Histograms along the axes show the distributions of eigenvector values.

**Extended Data Figure 9.**
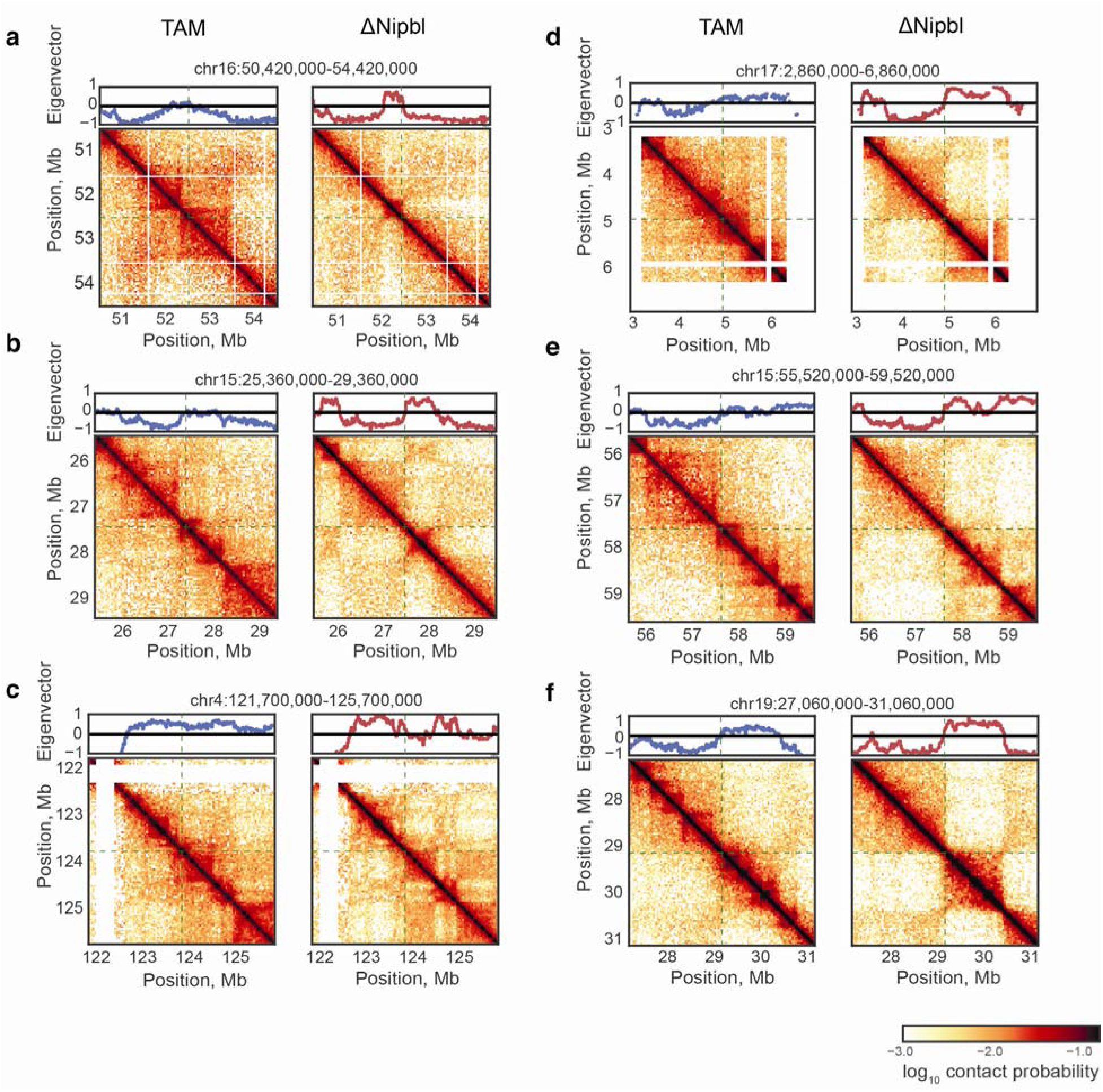
Boundaries of former TADs and new compartments do not coincide. Examples of TADs detected in WT cells which cross sharp compartment transitions revealed by ΔNipbl contact maps. Left column – TAM control data, right column – ΔNipbl data. Top of each figure – local eigenvector track in the corresponding cell type. The contact maps are centered at the sharp compartment transition in ΔNipbl. These examples illustrate that that chromatin-bound cohesins can locally interfere with genome compartmentalization.

**Extended Data Figure 10.**
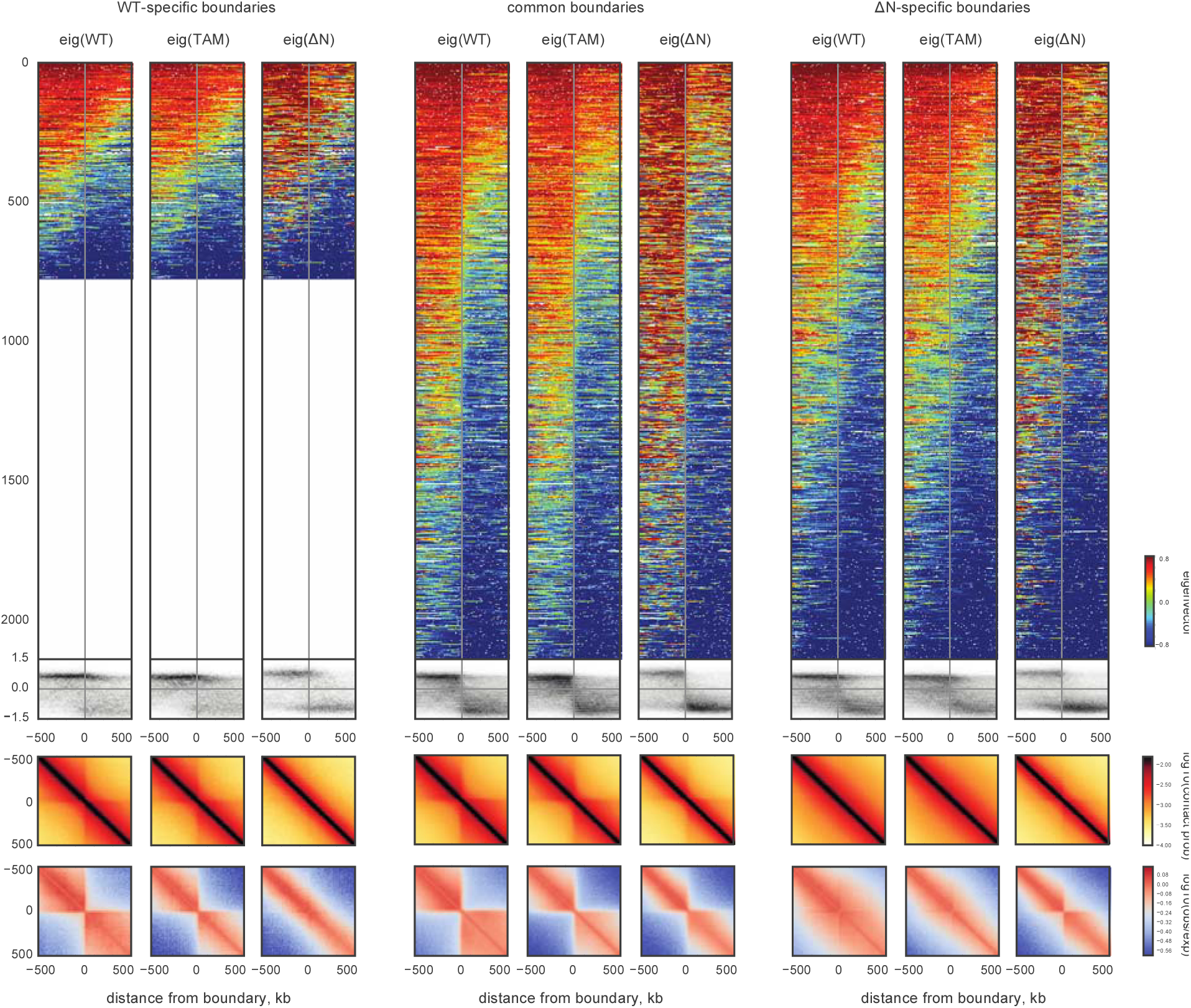
The residual contact-insulating boundaries in ΔNipbl are associated with compartment transitions. The first group of columns considers the boundaries detected in WT cells only, the second pair considers boundaries detected both in WT and ΔNipbl cells, the last pair considers boundaries detected in ΔNipbl only. For each group, the first, second and third columns display data (eigenvectors and Hi-C) from WT, TAM and ΔNipbl cells, respectively. Within each column: the top row – a stack of eigenvector tracks in a +/- 500kb window around boundaries, oriented such that the left-half of the window has greater average signal value and sorted by the average WT eigenvector value in the window. The second row – density histogram of eigenvector values as a function of the distance to the boundary. The third and fourth row – the boundary-centered average contact probability and observed/expected contact ratio, respectively. The density histograms show that common and ΔNipbl-specific boundaries correspond to sharp transitions of compartment signals in ΔNipbl cells, in contrast to the more diffuse signal at these positions in WT and TAM cells.

**Extended Data Figure 11.**
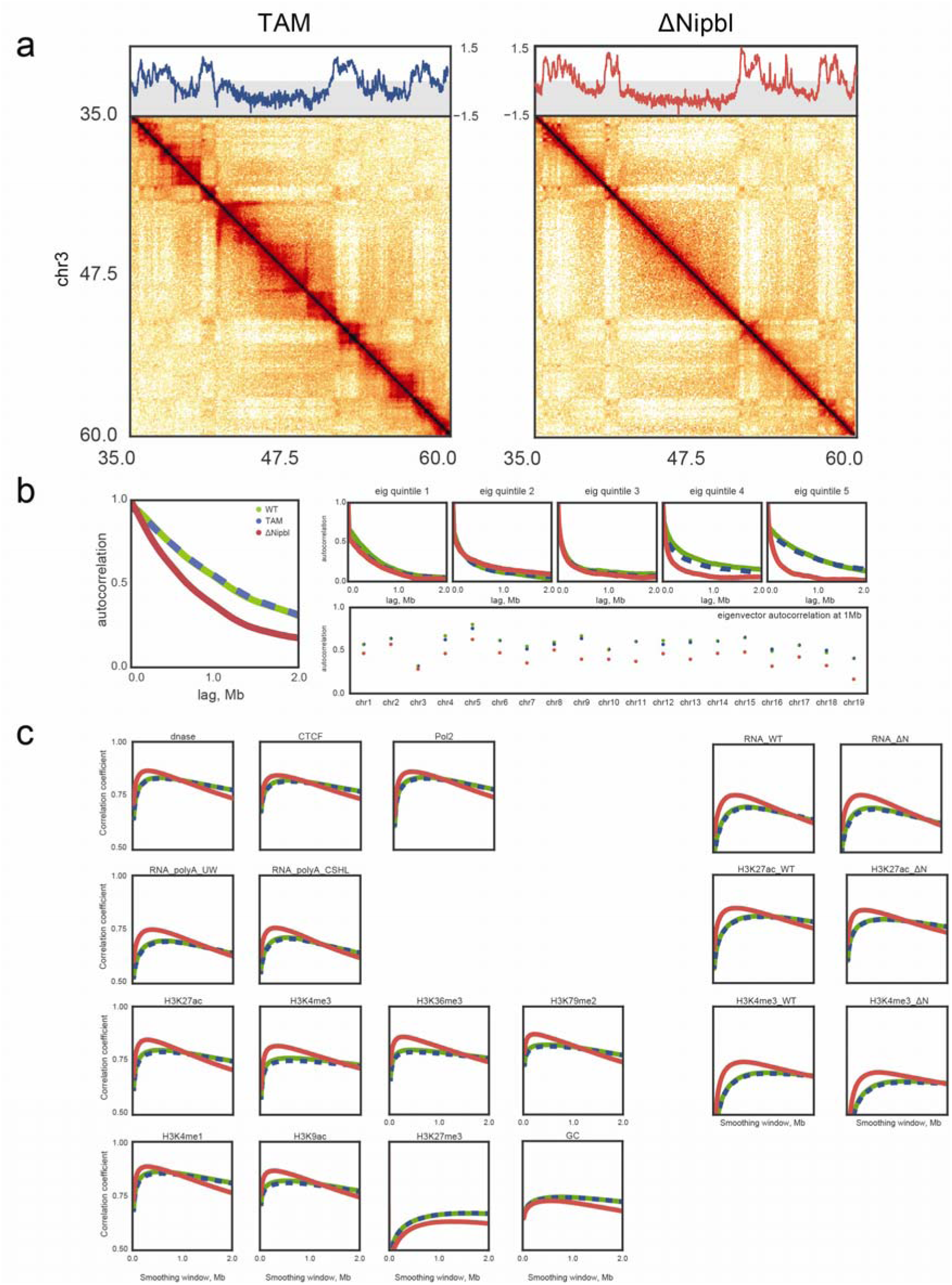
Fragmentation upon *Nipbl* deletion in smaller alternating regions of A and B compartment-type is activity-dependent. **(a)** Example region (chr3:35-60Mb) illustrating lack of compartment fragmentation in uniformly B-rich regions yet robust disappearance of TADs. Top – compartment eigenvector, Bottom – contact matrix snapshot. **(b)** Autocorrelation of eigenvector tracks reveals genome-wide fragmentation of active compartments. Left – the genome-wide correlation of the 20kb *cis* eigenvector values of pairs of loci as a function of their genomic separation (autocorrelation). Top right – eigenvector correlation of locus pairs split by quintile of the eigenvector value of the upstream locus. Bottom right – chromosome-wide values of eigenvector correlation of locus pairs separated by 1Mb. **(c)** Correlation between the smoothed histone and TF ChIP-seq and RNA-seq tracks and the 20kb *cis* eigenvectors as a function of the smoothing window size. Left group of panels – ENCODE data, right – data from this study. First and second rows – histone marks, third row – RNA-seq tracks, fourth row – miscellaneous tracks (DNase hypersensitivity, CTCF and PolII ChIP-seq and GC content). ΔNipbl eigenvectors show an increased correlation with tracks associated with transcriptional activity yet a decreased correlation with the repression-associated track of H3K27me3 and GC content.

**Extended Data Figure 12.**
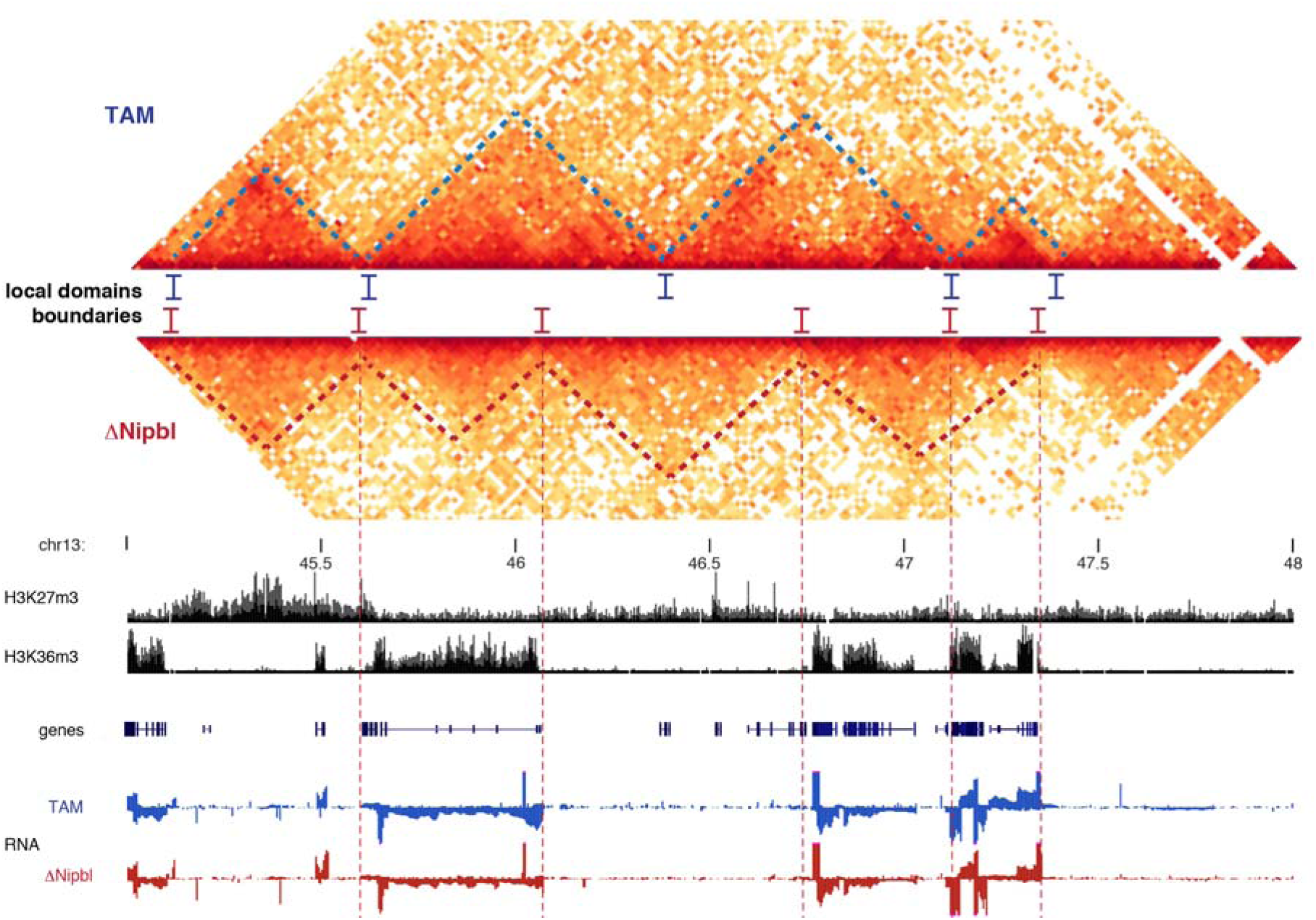
New structures appear in ΔNipbl cells and follow the underlying chromatin activity. Example of a large WT A-type compartment region (chr13:45-48Mb). Hi-C maps show different structures (TADs) highlighted by dashed lines (upper panel, TAM control, blue lines; lower panel, ΔNipbl, red line). Boundaries are shifted or lost and replaced by new ones in ΔNipbl cells. Histone ChIP-seq tracks ^42^ and stranded RNA-seq tracks (blue: TAM hepatocytes, red; ΔNipbl cells) highlight that WT/TAM TADs do not strictly follow the underlying chromatin activities, whereas the new structures in ΔNipbl cells delineated by red dashed lines correspond precisely to active versus inactive chromatin domains.

**Extended Data Figure 13.**
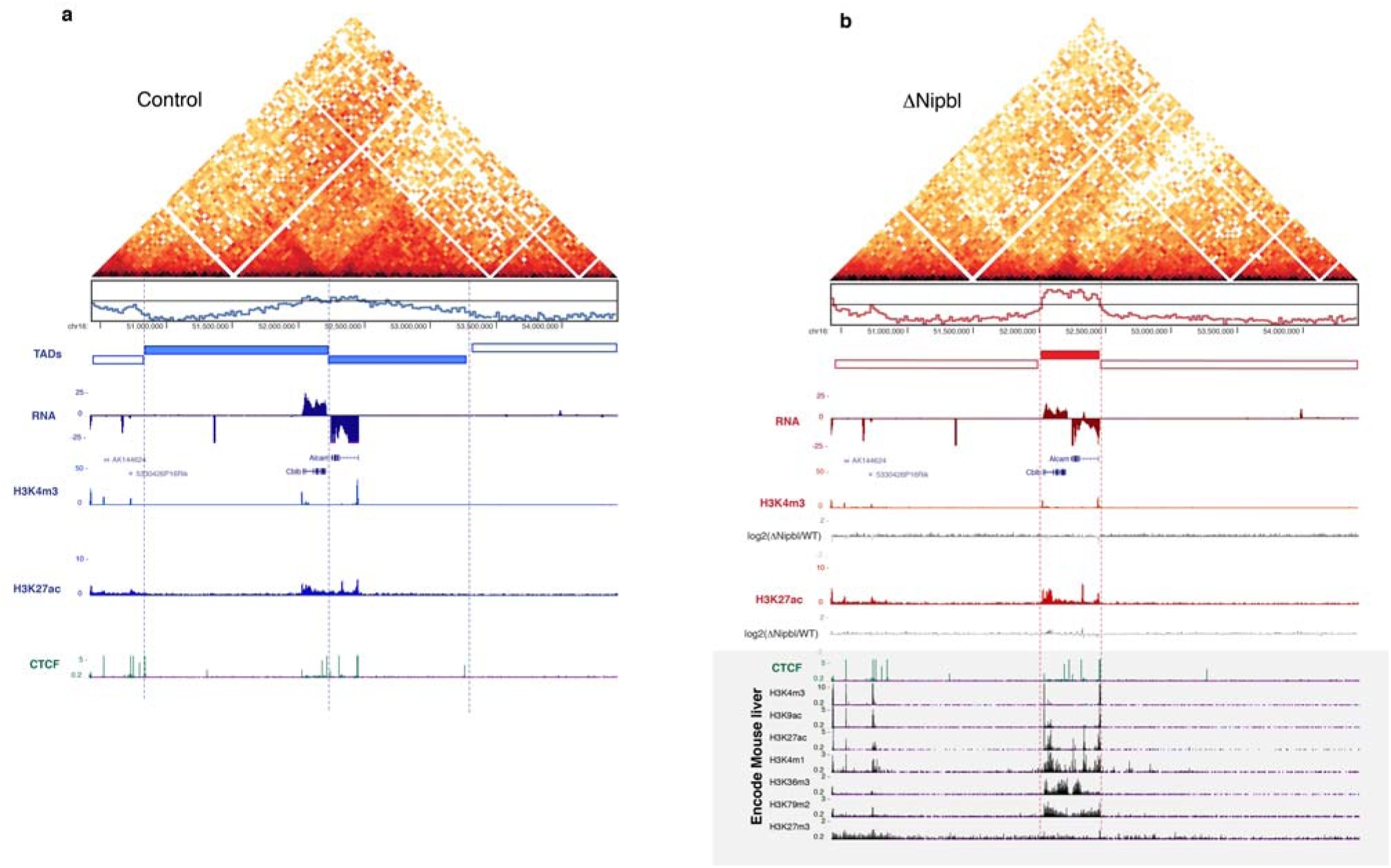
New compartments do not respect TAD boundaries but the underlying chromatin domains. A large region (chr16:50420000-54420000) adopts a very different 3D organisation in control **(a)** (in blue) and ΔNipbl cells **(b)** (in red). Hi-C data are shown, as well as the eigenvector values in the two conditions. RNA-Seq tracks showed minimal changes of expression (*Alcam* expression is reduced by 2-fold in ΔNipbl cells) and chromatin states. ChIP-Seq tracks for H3K27ac and H3K4me3 are shown in the two conditions, with log2 ratio tracks under the ΔNipbl **(b)** panel. Encode tracks (corresponding to WT liver cells) are shown in the grey boxed area. The new structure adopted in ΔNipbl cells put together the two active genes which are normally in different TADs in the same domain, corresponding to the active chromatin linear domain.

**Extended Data Figure 14.**
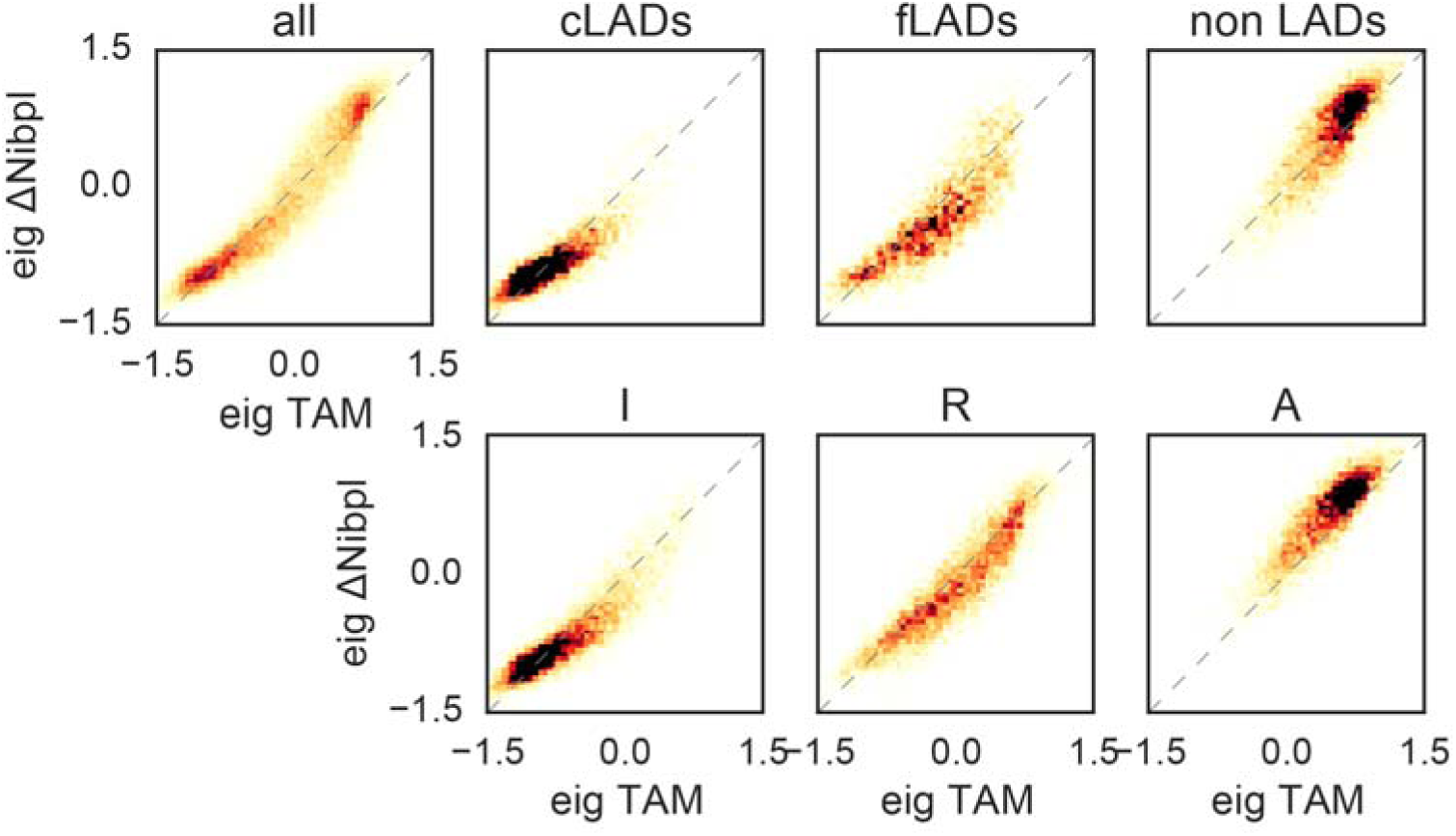
Correlation of *cis* eigenvector values of 100kb genomic bins before and after *Nipbl* deletion, split by the functional state of chromatin. Top row, left to right: genome-wide relationship; bins showing constitutive lamin-B1 association across 4 mouse cell types (cLADs); bins showing variable (facultative) lamin-B1 association (fLADs); binds not showing any association (non LADs). Bottom row: bins assigned the Inert ChromHMM simplified state; bins assigned the Repressed state; bins assigned the Active state.

**Extended Data Figure 15.**
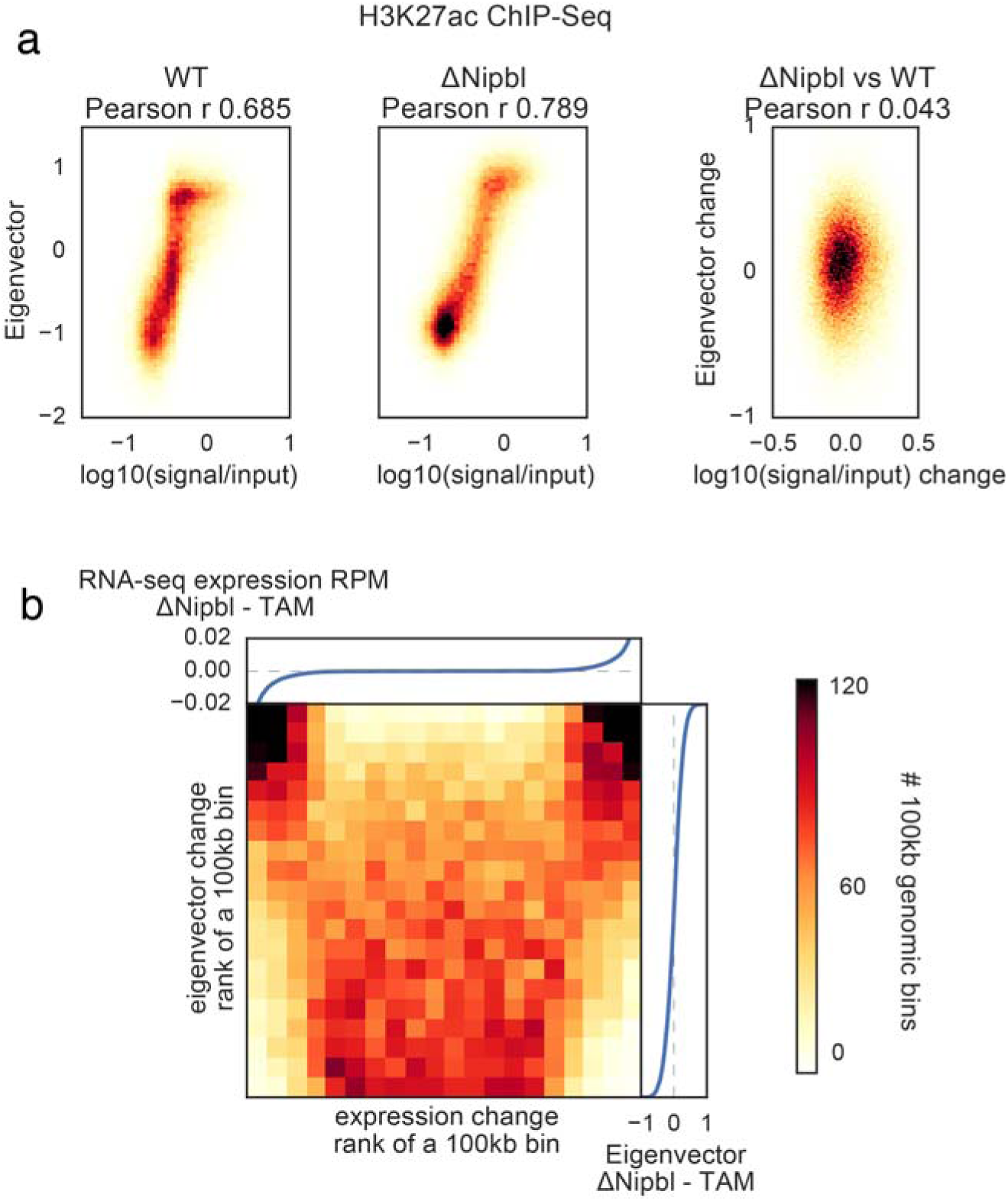
Eigenvector change upon *Nipbl* deletion is uncorrelated with changes in gene expression or epigenetic marks. **(a)** ChIP-Seq signal for histone marks of activity vs eigenvector value of 20kb bins, top row – H3K27ac, bottom row – H3K4me3. Left column – WT cells, middle column – ΔNipbl cells, right column – correlation of changes in both signals upon Nipbl deletion. **(b)** The change in the compartment structure upon *Nipbl* deletion cannot be attributed to the sign of the local expression change. The heatmap shows the number of 100kb genomic bins as a function of the ranks of expression change and the eigenvector change. The attached plots show the correspondence between the values of expression change (top) or eigenvector change (right) and their ranks.

**Extended Data Figure 16.**
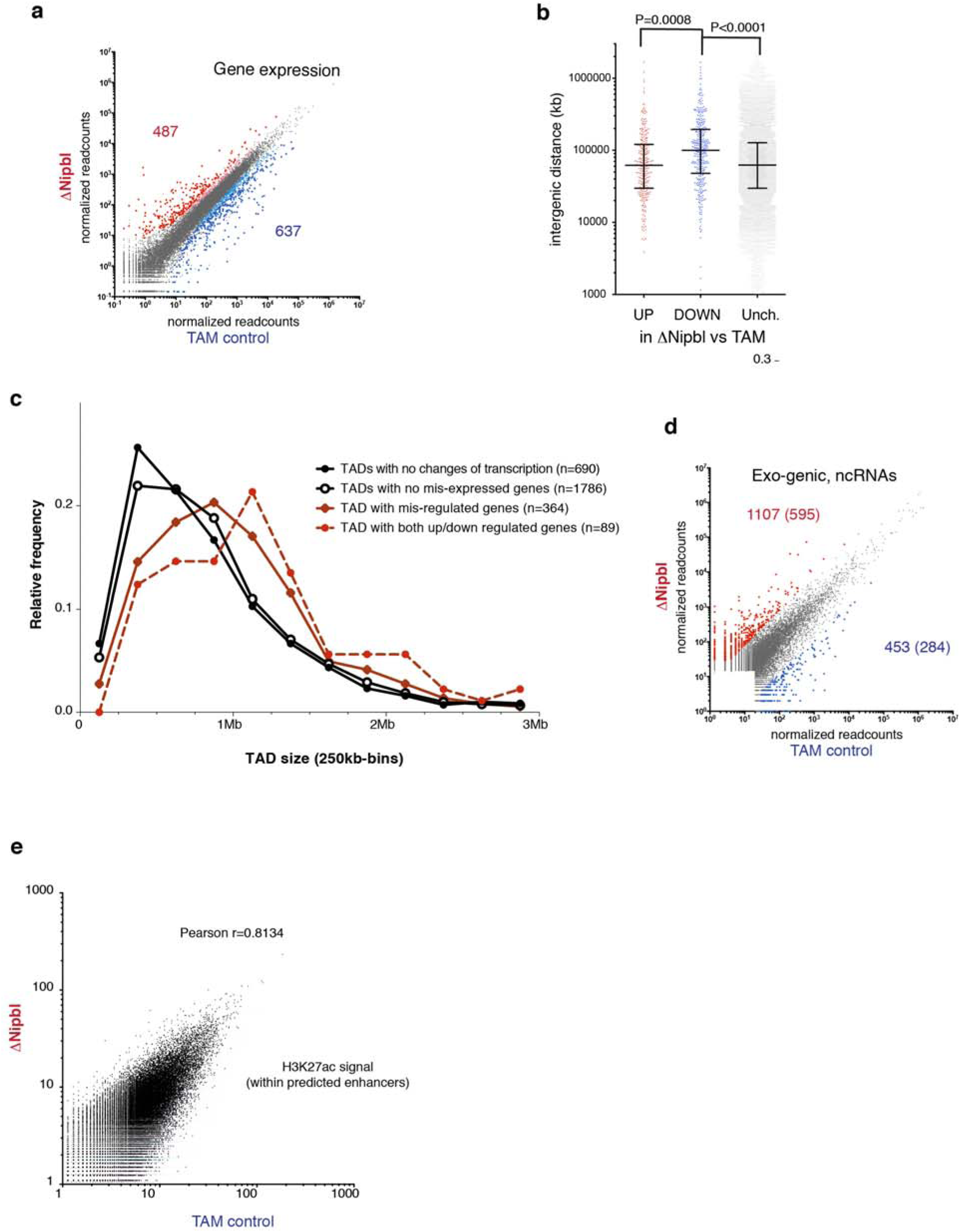
Expression changes in ΔNipbl hepatocytes. **(a)** Changes in gene expression between TAM controls and ΔNipbl liver cells (four replicates for each condition). Genes with significant changes in gene expression (FDR > 0.05) are coloured in red (upregulated) or blue (down-regulated), with larger dots corresponding to gene with a fold-change > 3 (numbers given correspond to these high-confidence subset of dis-regulated genes). **(b)** Intergenic distances for the different categories of dysregulated genes. Statistical differences determined by an unpaired two-tailed *t*-test. **(c)** Size distribution of the TADs observed in WT (lost in ΔNipbl) depending on the degree alteration of their transcriptional states. The size of TAD with transcriptional changes (red) is significantly larger than those that do not show transcriptional alterations (black) (Kolmogorov-Smirnov, *P*<0.0001) **(d)** Change in transcription in non-genic intervals (including inter-genic and antisense within gene bodies). Gene expression was calculated as the normalized number of read within intervals defined by merging adjacent 1kb windows showing readcounts over background (see *Methods*). The numbers of non-coding transcription up-regulated (in red) or down-regulated (in blue) in ΔNipbl compared to the TAM control is given (*P*-value <0.01, fold-change higher than 8), with the second number indicating the high-confidence events (labelled with coloured dots, expression value over an arbitrary threshold of 30 reads) which constitute the list used for subsequent analyses. **(e)** Comparison of control and ΔNipbl H3K27ac normalized signals within predicted liver enhancer elements (readcounts within +/- 500bp of predicted enhancer peak) ^42^.

**Extended Data Figure 17.**
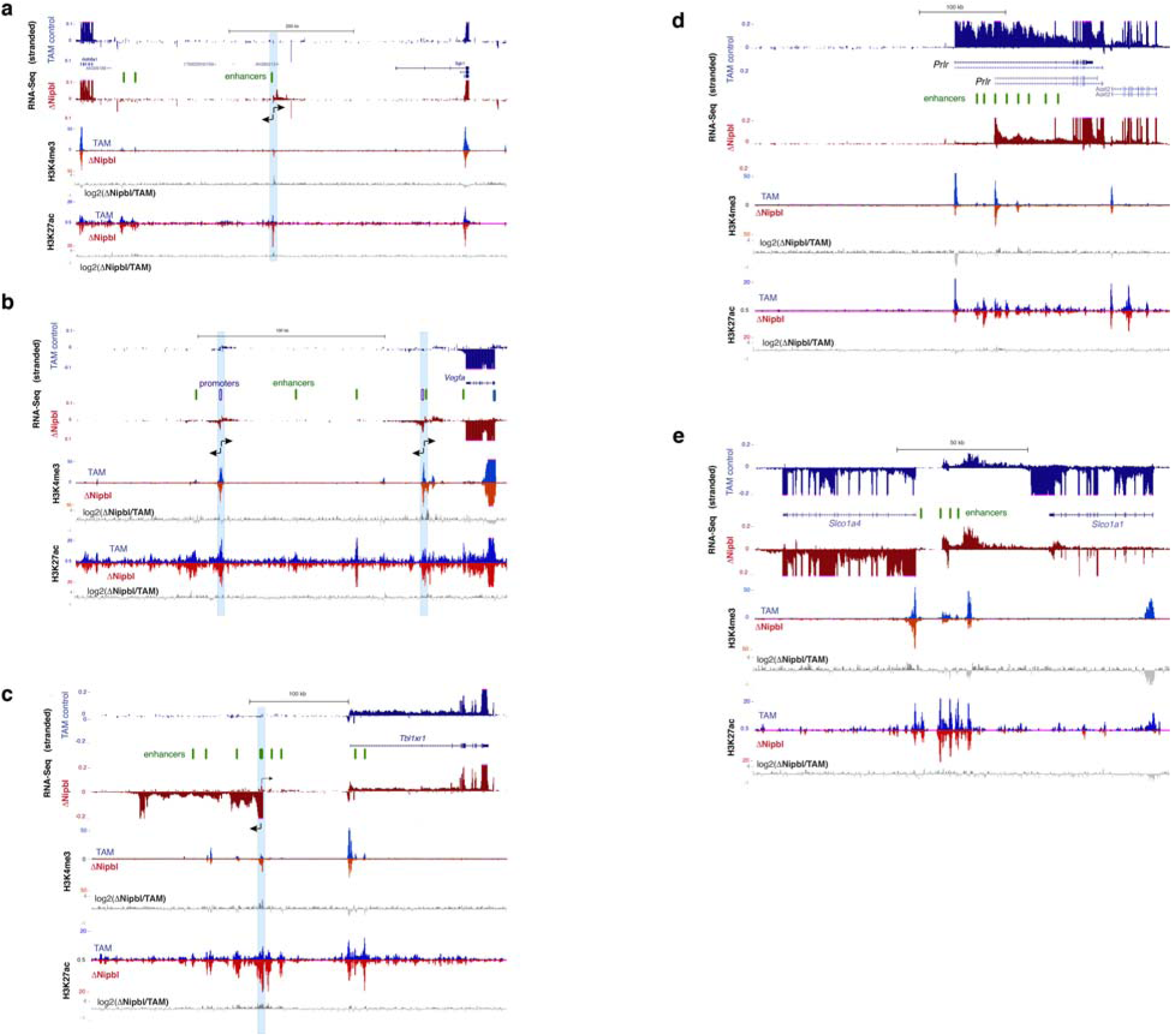
Transcriptional changes upon *Nipbl* deletion. Stranded RNA-Seq and ChIP-Seq tracks (H3K4me3, H3K27ac) are shown for control (blue) and ΔNipbl (red) samples. Comparison of the chromatin profiles are shown with log2(ΔNipbl/TAM) tracks for H3K4me3 and H3K27ac (in grey). Active enhancers (peaks of high H3K27ac, H3K4me1 ^42^, low H3K4me3) are depicted as green ovals. **(a)** chr10:21,090,000-21,781,000. Bidirectional transcription (position labeled with a blue bar) arises from an isolated enhancer in ΔNipbl cells. **(b)** chr17:45,945,000-46,176000. Bidirectional transcription (position labeled with a blue bar) arises from two cryptic promoters (H3K4me3 peaks, no/weak transcription in TAM) downstream of *Vegfa.* **(c)** chr3:21,712,500-22,126,240. A new transcript from a cryptic promoter 100 kb upstream of *Tbl1xr1*. H3K27ac signal is enhanced at peaks surrounding the activated cryptic promoter. **(d)** chr15:9,873,000-10,354,700. Promoter switch for *Prlr*, from an upstream promoter to a more downstream one surrounded by active enhancers. **(e)** chr6:141,743,961-141,904,692. Downregulation of *Slco1a1* and concomitant up-regulation of *Slco1a4* and non-coding intergenic transcripts (arrowheads). Distance of *Slco1a4* promoter to intergenic enhancers is less than 10kb, compared to 80 kb for *Slco1a1.*

**Extended Data Table 1.**
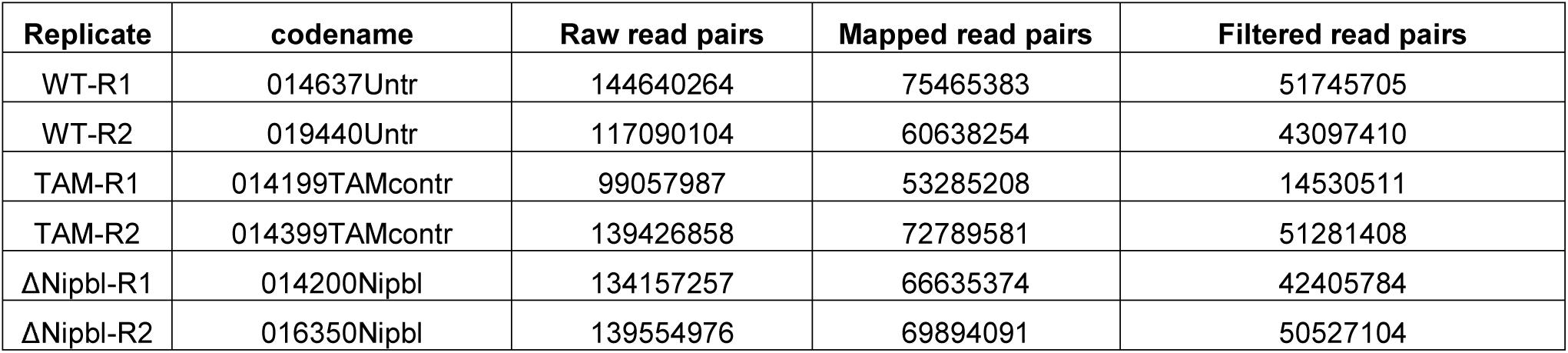
TCC libraries

**Extended Data Table 2.**
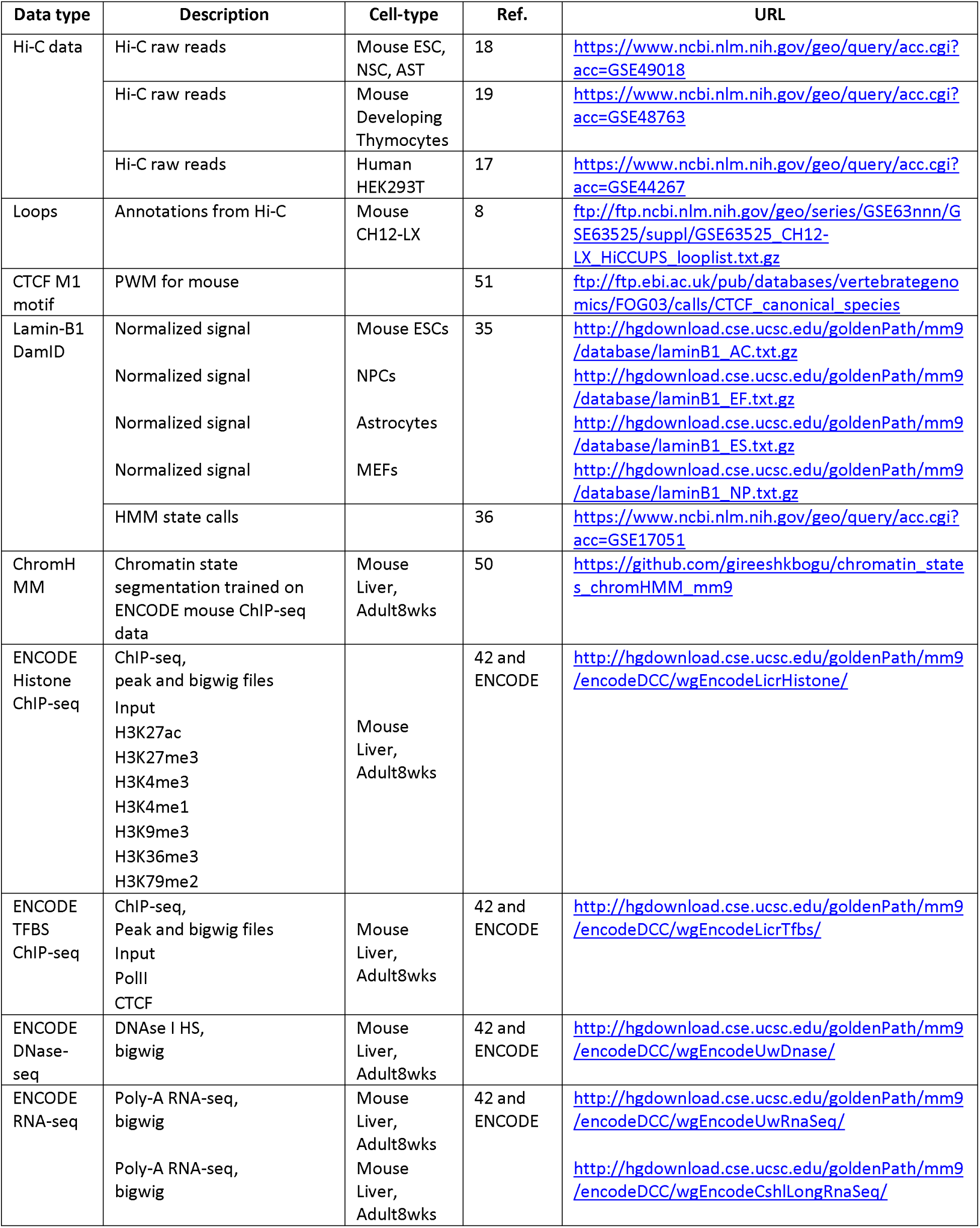
Public data sources

## Methods

### Experimental procedures

#### Generation of *Nipbl^flox/flox^* mice

The *Nipbl* locus was targeted by introduction of two *loxP* sites flanking exon 18 via homologous recombination in E14 mouse embryonic stem cells. Individual ESC clones were screened for successful recombination by Southern blotting deploying two unique probes hybridizing 5’ and 3’ to the integration site, respectively. Cells of a successful clone were injected into mouse blastocysts and resultant chimera were bred to C57BL/6J mice. Offspring were analyzed for successful germ line transmission by PCR and Southern blotting. *Nipbl^flox/+^* mice were maintained on C57BL/6J genetic background.

For deletion of the *floxed* exon, we used either a constitutive ubiquitous CRE-driver (Hprt::Cre ^39^), a limb-specific CRE-driver (Prx1::Cre ^40^) or an inducible, liver-specific *Cre* allele (*Ttr-cre/Esr1* ^43^.

Mice were genotyped by PCR using specifc primer pairs (details available on request).

All lines were maintained by breeding with C57Bl/6J mice. Mouse experiments were conducted in accordance with the principles and guidelines in place at European Molecular Biology Laboratory, as defined and overseen by its Institutional Animal Care and Use Committee, in accordance with the European Directive 2010/63/EU.

### Generation and preparation of *Nipbl^-/-^* adult primary hepatocytes

To inactivate *Nipbl* in adult mouse hepatocytes, *Nipbl^flox/+^* mice were crossed with mice carrying an inducible, liver-specific *Cre* allele (*Ttr-cre/Esr1* ^43^. Resultant double heterozygous mice were backcrossed to *Nipbl^flox/flox^*. For experiments we used animals homozygous for the floxed *Nipbl* allele either carrying one or no copy of the inducible, liver-specific *Cre* allele as sample (*Ttr-cre/Esr1^+/wt^*; *Nipbl^flox/flox^*) or control (*Ttr-cre/Esr1^wt/wt^*; *Nipbl^flox/flox^*), respectively.

12 week old mice were injected with 1mg Tamoxifen (100μl of 10mg/ml Tamoxifen in corn oil) on 5 consecutive days. After keeping these mice for another 5 days without injection, they were sacrificed and the hepatocytes were harvested. Until this time point, mice displayed no abnormal behavior, weight loss or any other obvious physiological changes. This was also the case, when we kept mice for additional 4 days without injection to test for any adverse effects immediately after our experimental time point.

The liver was dissected and the left lateral lobe was prepped for a two-step perfusion adapted from ^44,45^ First, the liver is perfused with an EDTA-containing buffer removing Ca^2+^ from the tissue. This weakens the integrity of the desmosome, which is then digested during the subsequent perfusion with a Ca^2+^ rich buffer containing collagenase. The freed hepatocytes were rinsed through a cell strainer and washed four times with ice-cold Ca^2+^ rich buffer without collagenase. For each wash the cells were spun at low centrifugal force (60g for 1 min), to reduce non-mesenchymal cells and debris, hence, enriching intact hepatocytes in the sample. Part of each sample was fixed with 1% PFA for 10 minutes at room temperature. Fixed and unfixed hepatocytes were aliquoted and frozen in liN_2_ for later use.

### *Nipbl* RNA levels and activity

Unfixed hepatocyte aliquots were thawed and RNA was prepared with Qiagen RNeasy Kit. cDNA was generated using NEB ProtoScript^®^ First Strand cDNA Synthesis Kit with random primer mix. RT-qPCR was performed with Applied SYBR Green PCR Master Mix and following primers: Nipbl-qPCR_F TCCCCAGTATGACCCTGTTT, Nipbl-qPCR_R AGAACATTTAGCCCGTTTGG, Gapdh-qPCR_F CTCCCACTCTTCCACCTTCG, Gapdh-qPCR_R CCACCACCCTGTTGCTGTAG, RTqPCR_Pgk1_Fwd TGGTATACCTGCTGGCTGGATGG and RTqPCR_Pgk1_Rev GACCCACAGCCTCGGCATATTTC.

For Western blots unfixed hepatocyte aliquots were lysed and fractionated with a Subcellular Protein Fractionation Kit (ThermoFisher). The blots were probed with antibodies against cohesin subunits SA-1 and SMC1 (a courtesy of Ana Losada, CNIO, Madrid) and Topo IIβ (611492, BD Biosciences) and Histone H2B (07-371, Millipore) as control for nuclear soluble and nuclear insoluble fractions, respectively.

### Immunohistochemistry on liver

Slices of adult livers were collected and fixed in 4% PFA overnight. After dehydration, the tissues were embedded in paraffin and sectioned at 6μm. The sections were deparaffinized with xylene, rehydrated and antigens were retrieved by boiling in citrate buffer. The sections were blocked in 10% FBS and incubated with primary antibodies (αphospho-Histone H3, 06-570 Millipore; αcleaved-caspase-3, #9661 Cell Signaling) at 4°C overnight. Primary antibodies were detected with goat anti-rabbit IgG Alexa Fluor^®^ 568 secondary antibody (A-11011, Invitrogen) and counter stained with DAPI. Images were acquired using confocal microscopy.

### RNA-seq libraries and sequencing

RNA integrity was tested with Bioanalyzer (Agilent RNA Nano Kit) and ribosomal RNA was removed using Ribo-Zero rRNA Removal Kit (Illumina) prior to library preparation. Strand-specific libraries were prepared with NEBNext^®^ Ultra™ Directional RNA Library Prep Kit for Illumina^®^. After amplification and size selection with Agencourt AMPure XP beads (Beckmann Coulter) their size-distributions were determined with Bioanalyzer. Equimolar pools of libraries were sequenced with Illumina HiSeq2000 (50bp, single end). We retrieved on average 25 mio reads per sample, of which 19 mio reads were uniquely mapped to the reference genome (NCBI37/mm9).

### ChIP-seq libraries and sequencing

Fixed aliquots of hepatocytes were hypotonically lysed and sonicated in 1% SDS/TE. An aliquot of each sample was reverse cross-linked in order to determine chromatin concentration and sonication efficiency. 20μg chromatin per sample was diluted in RIPA and incubated with 1.5μg of either αH3k4me3 antibody (C15410003-50, Diagenode) or αH3K27Ac antibody (ab4729, Abcam) at 4°C, overnight. The antibodies were retrieved with Dynabeads (IgA, Invitrogen) and bound chromatin was washed and eluted. After reverse cross-linking, the amount of ChIPped and input DNA was determined with Qubit (Thermo Fisher). The libraries were prepared with NEBNext^®^ ChIP-Seq Library Prep Kit for Illumina^®^. After amplification and size selection with E-Gel^®^ SizeSelect™ (Thermo Fisher) their size-distributions were determined with Bioanalyzer. Equimolar pools of libraries were sequenced with Illumina HiSeq2000 (50bp, single end). We retrieved on average 20 mio reads per sample, of which 15 mio reads were uniquely mapped to the reference genome (NCBI37/mm9).

### Tethered Chromatin Capture (TCC)

Roughly 100 mio fixed hepatocytes per sample were processed according to Kalhor et al.^30^ using HindIII. Libraries were PCR-amplified (12 cycles) and size selected with E-Gel^®^ SizeSelect™ (Thermo Fisher). Equimolar pools of libraries were sequenced with Illumina HiSeq2000 (50bp, paired end). We retrieved between 100 and 150 mio paired reads per sample, of which ~40% had both sides uniquely mapped to the reference genome (NCBI37/mm9).

## Computational analysis

### Preparation of Hi-C maps

We mapped the sequence of Hi-C molecules to reference mouse genome assembly mm9 using Bowtie 2.2.8 and the iterative mapping strategy, as described in ^46^ and implemented in the *hiclib* library for Python (publicly available at https://bitbucket.org/mirnylab/hiclib). Upon filtering PCR duplicates and reads mapped to multiple or zero locations, we aggregated the reads pairs into 20kb and 100kb genomic bins to produce Hi-C contact matrices. Low-coverage bins were then excluded from further analysis using the MAD-max (maximum allowed median absolute deviation) filter on genomic coverage, set to five median absolute deviations. To remove the short-range Hi-C artifacts - unligated and self-ligated Hi-C molecules - we removed contacts mapping to the same or adjacent genomic bins. The filtered 20kb and 100kb contacts matrices were then normalized using the iterative correction procedure (IC) ^46^, such that the genome-wide sum of contact probability for each row/column equals 1.0. Observed/expected contact maps were obtained by dividing each diagonal of a contact map by its average value over non-filtered genomic bins.

The same procedure was used to analyse other existing Hi-C datasets on cohesin-depleted cells ^17-19^.

### Compartment analysis via eigenvector decomposition

The compartment structure of Hi-C maps was detected using a modified procedure from ^46^. In short, compartments were quantified as the dominant eigenvector of the observed/expected 20kb and 100kb *cis* contacts maps upon subtraction of 1.0, as implemented in *hiclib*.

### Domain detection and loop coordinates

To identify contact domains, we used a segmentation algorithm very similar to ^47^, which divides the genome into domains in such a way as to maximize a global domain scoring function. We used two different scoring functions: one was the corner score function from ^8^ and the other was based on network modularity ^48^, which is a metric widely used to detect communities in networks. The modularity score for a domain spanning genomic bins *a* to *b* inclusively is given by

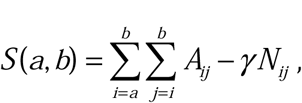

where *A* is the contact matrix and *N* is the corresponding matrix of a penalizing background model. The resolution parameter controls the strength of the penalty and therefore the characteristic size of the domains identified.

By restricting the solution space to contiguous segmentations, both calculating domain scores and finding the highest scoring segmentation can be reduced to O(n^2) dynamic programming algorithms. Optimal segmentation, in particular, becomes the well-known max-sum algorithm on a weighted directed acyclic graph ^49^. Furthermore, one can marginalize over the space of all possible segmentations to obtain a linear genomic track of boundary scores for a given domain scoring function, also in O(n^2) time. The implementation of these and related algorithms is provided in the lavaburst package (https://github.com/nezar-compbio/lavaburst).

The domain calls used in composite heatmaps were computed using the corner score on 20kb resolution matrices. To robustly call insulating boundaries across different conditions, we exploit the multi-resolution nature of the modularity score and compute the average marginal boundary scores on 20kb WT and ΔNipbl contact matrices sweeping over a range of gamma values to obtain a 1D boundary (i.e., insulation) track. Short intervals representing insulating loci were called by thresholding on the boundary score, and the common and unique loci to each condition were determined by interval intersection.

To characterize the structure of known loops in our data, we used the list of loops detected in Hi-C maps for CH12-LX mouse cell line in ^8^.

### Analysis of epigenetic features

To study the influence of the epigenetic chromatin state on genomic architecture, we used the ChromHMM genomic state annotation from ^50^. Briefly, the authors trained the ChromHMM genomic segmentation model on H3K4me1, H3K4me3, H3K36me3, H3K27me3, H3K27ac, CTCF and RNA polymerase II ENCODE tracks for mouse liver. This model assigned to each genomic loci one of 15 possible epigenetic states. For our analyses, we further grouped these 15 states into three: Active (states 1-11, 14 and 15 characterized by presence of PolII), Repressed (states 10-12, characterized by high H3K27me3 and low H3K27ac emission probabilities) and Inert (state 13, lack of any signal).

To find the average footprint of a chromatin-bound CTCF, we detected all occurrences of the M1 CTCF motif^51^ in the mm9 genome (filtered by p-value > 0.0001), intersected with CTCF binding peaks in ENCODE adult mouse liver dataset^52^. This procedure yielded 27840 peaks.

To study how lamina association affects genome compartmentalization, we used the dataset of Lamin-B1-binding loci from ^35^ containing data from four mouse cell lines: embryonic stem cells (ESC), multipotent neural precursor cells (NPC), terminally differentiated astrocytes (AC) and embryonic fibroblasts (MEF). We then selected 100kb genomic bins with more than 60 locations probed for lamin-B1 binding. We called the bins as constitutive LAD (cLAD) or non-LAD bins if >90% of probed locations were bound to lamina or not bound to lamina across all four tested cell lines, respectively. Finally, bins were called facultative LADs if for more than 90% of probes showed lamina binding in some cell lines, but not in others.

### RNA-Seq

We mapped the RNA-seq data to mm9 reference mouse genome assembly and GENCODE vM1 transcriptome ^53^ using STAR v.2.5.0a ^54^ and scripts from the ENCODE RNA-Seq pipeline [https://github.com/ENCODE-DCC/chip-seq-pipeline]. To obtain the tracks of local transcription, we aggregated the uniquely mapped reads into RPM-normalized bigWig files using the built-in STAR functionality. To find differentially expressed genes, we aggregated the read counts at the gene level using HTSeq ^55^ with the "union” option and called DE genes with DESeq2 ^56^.

For identification of exogenic transcripts, we aligned reads on the mm9 genome using HISAT2^57^ with the option to authorize splicing events. From these alignments, we count the number of reads in sliding windows of 1kb (with a step of 600nt) taking each strand of the genome as separate. We use a gaussian mixture model (2 distributions) to find a cutoff value for readcounts differentiating "expressed” regions from noise. We choose a cutoff of 15 reads according to the best fitted gaussian distributions. To avoid overestimating the number of exogenic transcripts, we merged adjacent expressed windows (on the same strand) as composite transcripts. We used spliced reads to merge expressed windows that were not directly adjacent, but linked by spliced reads, with a cutoff of 7 splice reads. The resulting merged and linked windows defined the region corresponding to the exogenic transcription unit, and we took its 5’ end (using the strand information, and after trimming the 5’end of all nucleotides with no coverage) as transcriptional start site.

### ChIP-Seq

We mapped the sequences obtained in ChIP-seq experiments following the ENCODE 3 pipeline [https://github.com/ENCODE-DCC/chip-seq-pipeline].

### Simulations of loop extrusion

To investigate the impact of NIPBL depletion on TADs and loops in the context of loop extrusion, we performed simulations that couple the 1D dynamics of loop extrusion by LEFs with 3D polymer dynamics, as previously^21^. We then generated contact maps and calculated P(s), also as previously described.

We considered a system of 42 consecutive TADs of 400kb, where impermeable BEs were placed between each pair of neighboring TADs. We modeled the chromatin fiber as a series of 1kb monomers (~6 nucleosomes, ~20nm), such that each TAD was 400 monomers. Polymer connectivity, stiffness, and excluded volume were implemented as previously described ^21^. Extruded loops held by LEFs were implemented by connecting the two monomers held by the two heads of each LEF with a harmonic bond, as previously^21,58^.

Three-dimensional polymer dynamics were implemented using OpenMM, a high-performance GPU-assisted molecular dynamics API ^59,60^. Simulations were initialized from a compact polymer conformation, as described in [PMID: 25472862], created on a cubic lattice a box of the size (PBC box – 2). Prior to a block of simulations, LEF dynamics were advanced by 500,000 steps. To allow the polymer fiber to equilibrate with this set of LEF-imposed bonds, simulations were then advanced for 400 blocks of simulations. After that, 2000 blocks (ie. blocks of 1D+3D dynamics) of simulations were performed and their conformations were recorded. After that, LEF dynamics were advanced by 500,000 steps, and the process was repeated, until 5000 conformations for each parameter set were obtained.

For the control simulation, we considered LEFs with 200kb processivity and 400kb separation. To investigate the effect of depleting the amount of bound cohesin, we then increased LEF separation by 2-fold, 4-fold, and 8-fold. All simulations used a stiffness of 2, density of 0.2, 3D-to-1D-steps of 2500. To generate contact maps from simulated conformations, a capture radius of 6 was used.

For display, simulated contact maps were first binned using 10 monomer (40kb) bins. Then, the map was normalized such the average value of the first diagonal equals one, and log10 transformed. Finally, the color-scale was clipped to show 1.5 logarithmic orders of magnitude as its dynamic range.

To calculate *P(s)* plots from simulated data, a contact map with 1 monomer resolution was used. For each diagonal of the contact map, we determined which regions within 400-monomer TADs and which were between TADs. We then averaged the values in logarithmically-spaced bins of increasing distance with a step of 1.1. Similar to experimental data, P(s) curves were then vertically shifted such that P(s) at 10kb was equal to 1.

